# Chromosome folding encodes mechanisms of cohesion maintenance

**DOI:** 10.64898/2026.11.02.763844

**Authors:** Xingchi Yan, Maxime M.C. Tortora, Anton Goloborodko, Geoffrey Fudenberg

## Abstract

After DNA replication, sister chromatids are physically linked by the cohesin complex. These cohesin linkages must be maintained for many hours for faithful chromosome segregation and DNA repair. Cohesin interacts with multiple rapidly exchanging cofactors, including SORORIN, which is often described as a ‘molecular lock’ for cohesin on chromatin. How such transient SORORIN binding can protect cohesin over much longer timescales remains mechanistically unclear. Furthermore, how loop-extruding cohesins interact with and influence the positioning of sister-linking cohesins remains largely unknown. Here we show that changes in sister chromatid folding over time and after perturbations encode distinct signatures of the underlying mechanisms of cohesion maintenance. By integrating biochemical kinetics with polymer modeling, we infer cohesin turnover, motion, and encounter rules from these folding signatures. Repeated transient sequestration by SORORIN enables long-lasting protection yet acts as a quantitative modulator, rather than a binary switch for cohesion maintenance. Prolonging cohesin residence time by WAPL depletion exposes additional mechanisms shaping sister chromatid conformation, including pushing of sister-linking cohesins and diffusion of loop-extruding cohesins. Together, these results reveal how transient cofactor interactions regulate long-lived cohesin maintenance and establish sister chromosome organization as a quantitative readout of otherwise hidden molecular mechanisms.

## Introduction

Sister chromatid cohesion is established in S-phase alongside DNA replication, maintained throughout G_2_, and ultimately released to permit chromosome segregation (**Fig. 1a**) (*1*, *2*). This long-lived physical linkage depends on cohesin complexes that are stably associated with chromatin (*3*). Stable cohesin binding is typically attributed to acetylation and subsequent protection by the cofactor SORORIN (*4–6*). SORORIN depletion (ΔSORORIN) leads to insufficient cohesion and premature sister chromatid separation in mitotic spreads. This strong phenotype, together with structural and biochemical evidence (*5*, *7*, *8*), has motivated a view that SORORIN “locks” cohesin on chromatin, functioning as a ‘molecular switch’ for cohesion (*1*, *9*). Conversely, depletions of the cofactors WAPL (ΔWAPL) or PDS5 display over-cohesion, where arms of sister chromatids adopt a ‘zipped’ morphology in mitosis (*4–6*). Yet these cofactors interact with cohesin only transiently (*10*), leaving unresolved how transient interactions sustain long-lived cohesion and organize replicated chromosomes.

**Figure 1.**
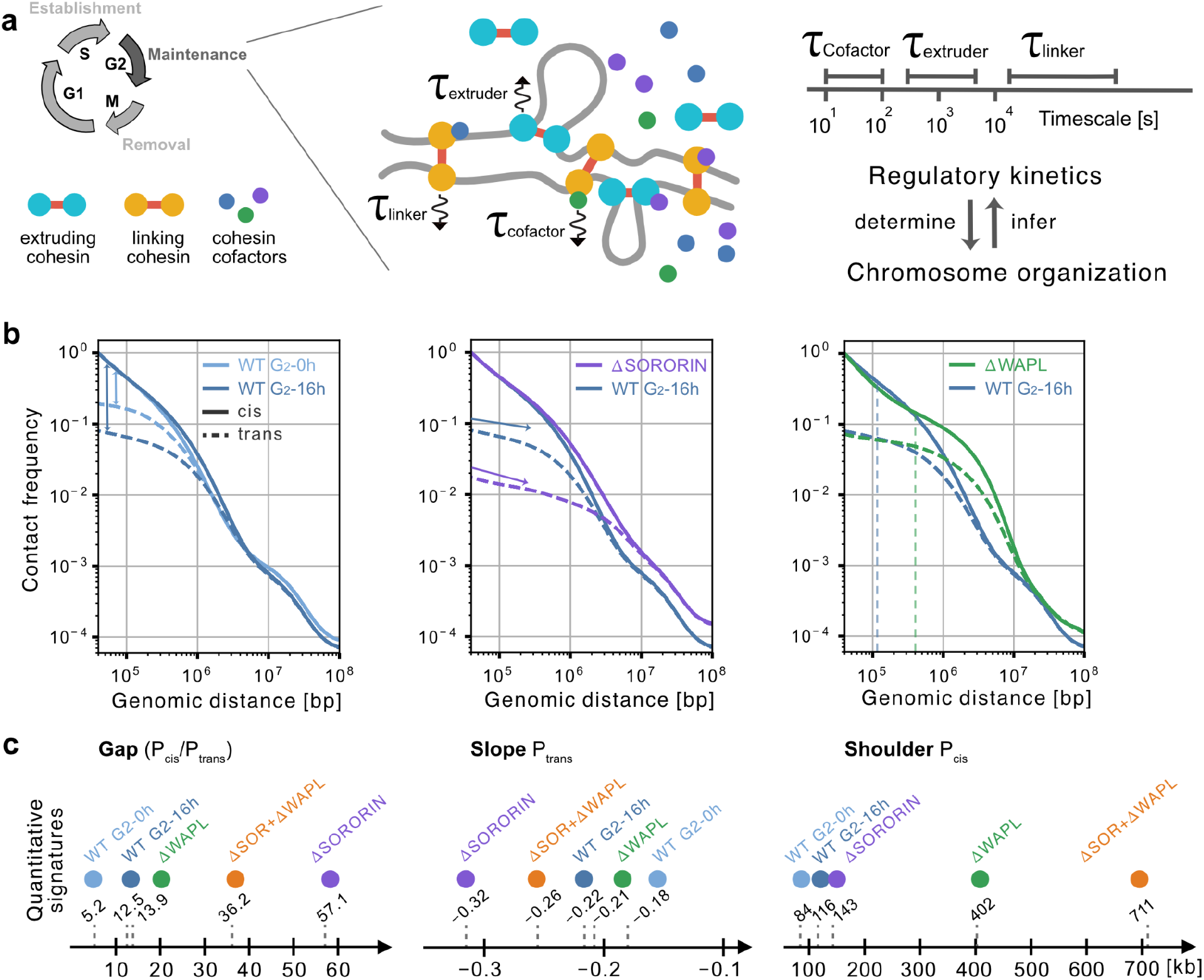
Regulatory kinetics of sister-linking and loop-extruding cohesins determine sister chromatid organization. **(a)** Illustration of cohesin dynamics and characteristic timescales through the cell cycle. After cohesion establishment in S phase, sister chromatid cohesion is maintained throughout G_2_. Sister-linking cohesins persist for hours, whereas loop-extruding cohesins and cofactors exchange on much shorter timescales. **(b)** Sister-chromatid-sensitive Hi-C (scsHi-C) contact-frequency curves for within-chromatid contacts (P_cis_, solid) and between-sister contacts (P_trans_, dashed) for: time in G_2_ (left) and after SORORIN depletion (ΔSORORIN; middle) (*21*), and after WAPL depletion (ΔWAPL; right) (*6*). Lines showing the gap (vertical arrows), the slope (tangent arrows), and the shoulder position (vertical dashes) are overlaid on respective panels. **(c)** Quantification of three signatures: the gap, defined as the ratio between P_cis_ and P_trans_ at 40 kb (left); the slope of P_trans_ quantified at 40 kb (middle); and the position of the P_cis_ shoulder (right). WT G_2_ progression alters the slope and gap, ΔSORORIN greatly increases the gap, and ΔWAPL shifts the shoulder of P_cis_ .

Mechanistic understanding remains experimentally limited because the molecular processes governing cohesion maintenance cannot be directly observed *in vivo*. First, cohesion has traditionally been assessed using mitotic spreads, which distinguish intact from defective cohesion at a single time point. Such experiments do not measure how cohesion changes over time, despite evidence that it is continuously regulated and sister chromatids slowly separate throughout G_2_ (*11–13*). Second, cohesin can either extrude chromatin loops (***extruders***) or link sister chromatids (***linkers***), yet both populations share a common pool of regulators (**Fig. 1a**). Because these populations are difficult to distinguish *in vivo*, the effects of individual regulators on each population cannot readily be disentangled. Third, how encounters between linkers and extruders are resolved may shape sister chromosome organization. *In vitro* observations of cohesin encounters revealed that blocking, bypass, pushing, and cohesin removal can all occur (*14*). This motivates several possible rules for extruder-linker encounters *in vivo* (*15–18*), yet which of these rules operate remains unclear. Such mechanistic understanding will ultimately be required to predict how perturbations to cohesin regulation reshape sister chromosome organization and downstream processes, including homology-directed DNA repair (*19*, *20*).

Although the molecular interplay that maintains cohesion cannot currently be directly observed in living cells, it nevertheless shapes sister chromosome folding. Emerging sister-resolved chromosome conformation capture (*6*, *21–23*) (e.g. scsHi-C (*6*, *21*, *22*)), combined with degron-based cohesin perturbations (*24*) and *in silico* polymer models (*25*), provide an opportunity to infer key aspects of this hidden molecular interplay. Polymer models have previously shown that chromosome conformation can be used to infer extruder velocity and residence time in interphase (*26–28*), rules of engagement between condensins and cohesin in mitosis (*16*), and sister chromatid misalignment after replication (*17*, *29*). Chemical reaction networks can further relate cofactor abundance to cohesin kinetics (*28*). Still, the respective contributions of extruders, linkers, and their regulators to sister chromosome folding during cohesion maintenance remain unresolved.

Here we show that the organization of sister chromosomes encodes quantitative information about the regulatory kinetics and rules of engagement of cohesin. To decode this information, we developed an integrated polymer model and reaction network framework. Applying our framework to time-resolved scsHi-C in wild-type cells excludes both rapidly exchanging and permanent linkers, identifying a residence time of ∼11 h for sister-linking cohesins. This long residence time arises from repeated SORORIN-dependent sequestration, which protects linkers from unloading. WAPL depletion exposes additional contributions from extruder diffusion and extruder-linker pushing that are largely masked in wild-type cells. Together, these results establish that mechanisms underlying cohesion maintenance can be inferred quantitatively from sister chromosome folding.

## Results

### Sister chromosome conformation changes with time in G_2_ and cofactor abundance

Contact versus distance curves, or P(s), provide informative summaries of genome-wide folding (*11*, *12*, *30*). For conventional Hi-C, a key feature of these curves is the position of the shoulder, which relates approximately to extruded loop size (*30*, *31*) and can be quantified as the genomic separation, s, at which the derivative of P(s) is maximal (s[max(P’)]). For scsHi-C, which generates curves both within a chromosome (P_cis_) and between sister chromatids (P_trans_, **Fig. 1b**), analogous features have been less thoroughly characterized. From experimental data, we identified three features of these curves: (i) the gap, or ratio between P_cis_ and P_trans_; (ii) the slope of P_trans_; and (iii) the position of the P_cis_ shoulder, defined analogously to the feature in conventional Hi-C curves. These features evolve over time in G_2_ and are altered by perturbations to cohesin cofactors. Progression through G_2_ decreases the P_trans_ slope and increases the gap; ΔSORORIN increases the gap, whereas ΔWAPL shifts the P_cis_ shoulder toward larger genomic distances (**Fig. 1b,c**). Which mechanisms contribute to these differences, and how they relate quantitatively to cohesin molecular kinetics remain unknown. We thus first used mechanism-agnostic polymer simulations to ask how properties of linker and extruder positioning influence these folding signatures, and then turned to mechanistic simulations to infer cohesin kinetics and encounter rules from experimental data.

### Polymer simulations map cohesin linker and extruder positions to 3D conformation

To determine how cohesin positioning gives rise to these distinct folding signatures, we developed mechanism-agnostic polymer simulations of sister chromatids at 2.5 kb resolution organized by linkers and extruders (**Fig. 2a**, **Methods**). Following previous work (26), linkers connected one position on each sister chromatid, whereas extruders connected two positions along a single chromatid (15). We then varied the positioning of linkers and extruders in simulations, generated ensembles of conformations, and computed P_cis_ and P_trans_ curves.

**Figure 2.**
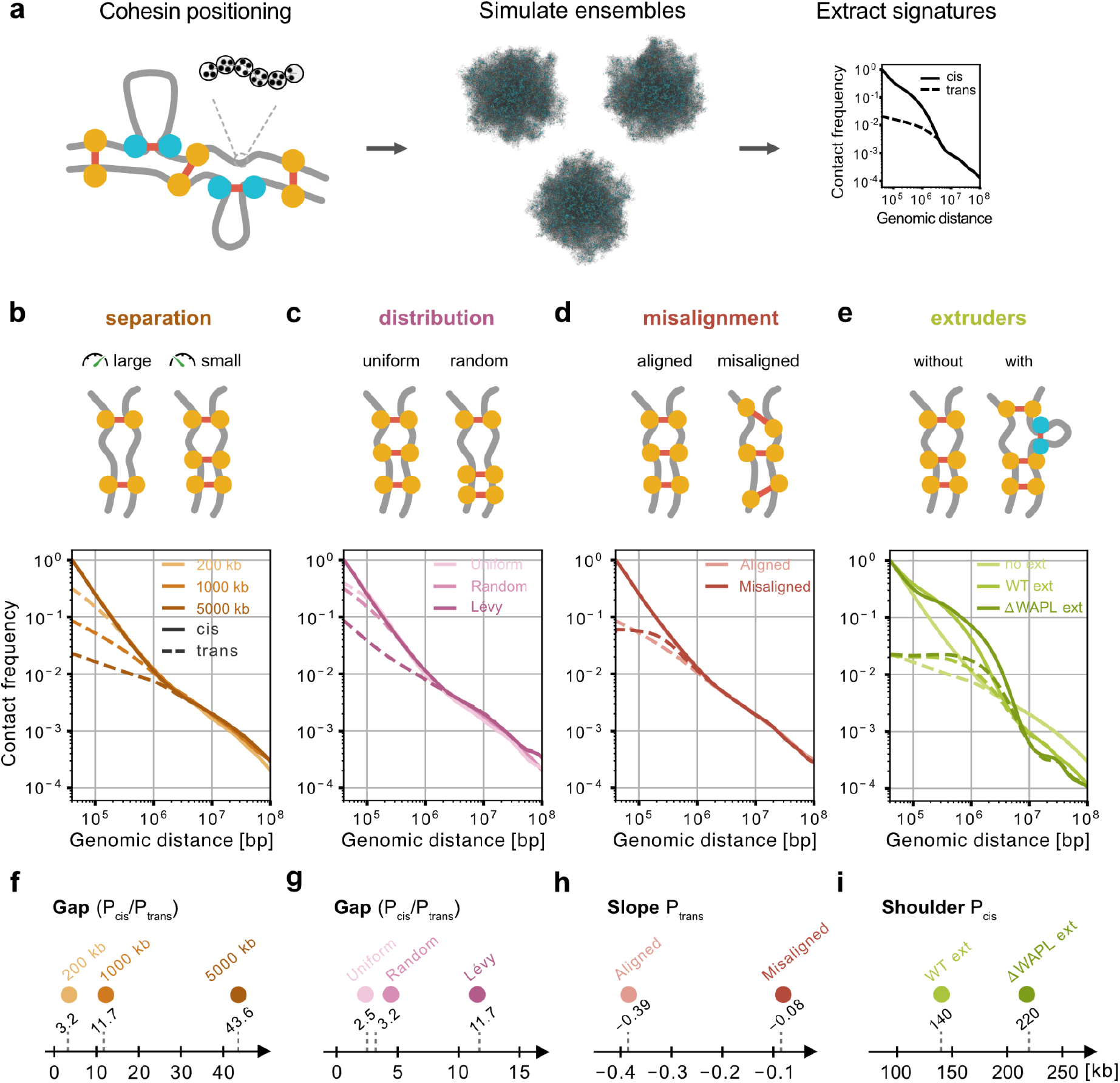
Polymer models map cohesin positioning to 3D genome organization within and between sister chromatids. **(a)** Schematic of how polymer simulations map cohesin linker and extruder positions to contact-frequency curves. (**b-e)** Simulations were used to test the following factors for their influence on 3D genome organization: **(b)** separation between sister linkers; **(c)** the degree of clustering of sister linkers along the genome (at fixed 200 kb separation); **(d)** the misalignment of sister linkers (at fixed 1 Mb separation); and **(e)** the presence of extruders (at fixed 5 Mb separation). Plots below each illustration show corresponding contact-frequency versus distance scaling curves, with P_cis_ within chromatids (solid lines) and P_trans_ between chromatids (dashed lines). **(f–i)** Quantification of the corresponding signatures of sister chromosome organization. **(f–g)** the P_cis_/P_trans_ gap for varying linker separation **(f)** and linker overdispersion **(g)**. **(h)** The slope of P_trans_ for varying linker misalignment. **(i)** The P_cis_ shoulder for varying extruder kinetics.

In simulations, the average separation between linkers primarily controlled the gap between P_cis_ and P_trans_, but also modulated the P_trans_ slope (**Fig. 2b,c, Fig. S1a**). Misalignment, defined for each linker as the genomic separation between anchor positions on each sister, primarily controlled the P_trans_ slope (**Fig. 2d, Fig. S1b,c**). Finally, the addition of extruders strongly altered P_cis_, with smaller effects on the P_trans_ slope and gap (**Fig. 2e**). Thus, P_cis_ and P_trans_ encode distinct information about linker separation, misalignment, and extruded loop size. The increased gap from G_2_ entry to 16 h indicates an increased effective linker separation during G_2_ (**Figs. 1b, 2b,c**) and argues against a simple model in which cohesion is permanently locked from S phase until mitosis. We therefore developed mechanistic simulations that jointly model extruders, linkers and their regulation by cofactors to determine what timescale of linker turnover could account for cohesion maintenance.

### The ultra-long timescale of cohesion maintenance can be inferred from scsHi-C

To infer the timescale of unloading *in vivo* and how it is regulated by cofactors, we performed mechanistic simulations that included extruder, linker and cofactor dynamics (**Fig. 3a; Fig. S3a, b**). Initial linker misalignment and separation were determined by scsHi-C at G_2_ entry (∼141 kb and 275 kb, respectively, **Fig. S2b-d**). We modeled cofactor interactions with both extruders and linkers by extending the chemical reaction network approach developed previously for extrusion (*28*) (**Fig. S3a, b, Methods**). Specifically, linkers could reversibly bind SORORIN and PDS5 before undergoing irreversible WAPL-mediated unloading (**Fig. 3a**; **Fig. S3a,b**).

**Figure 3.**
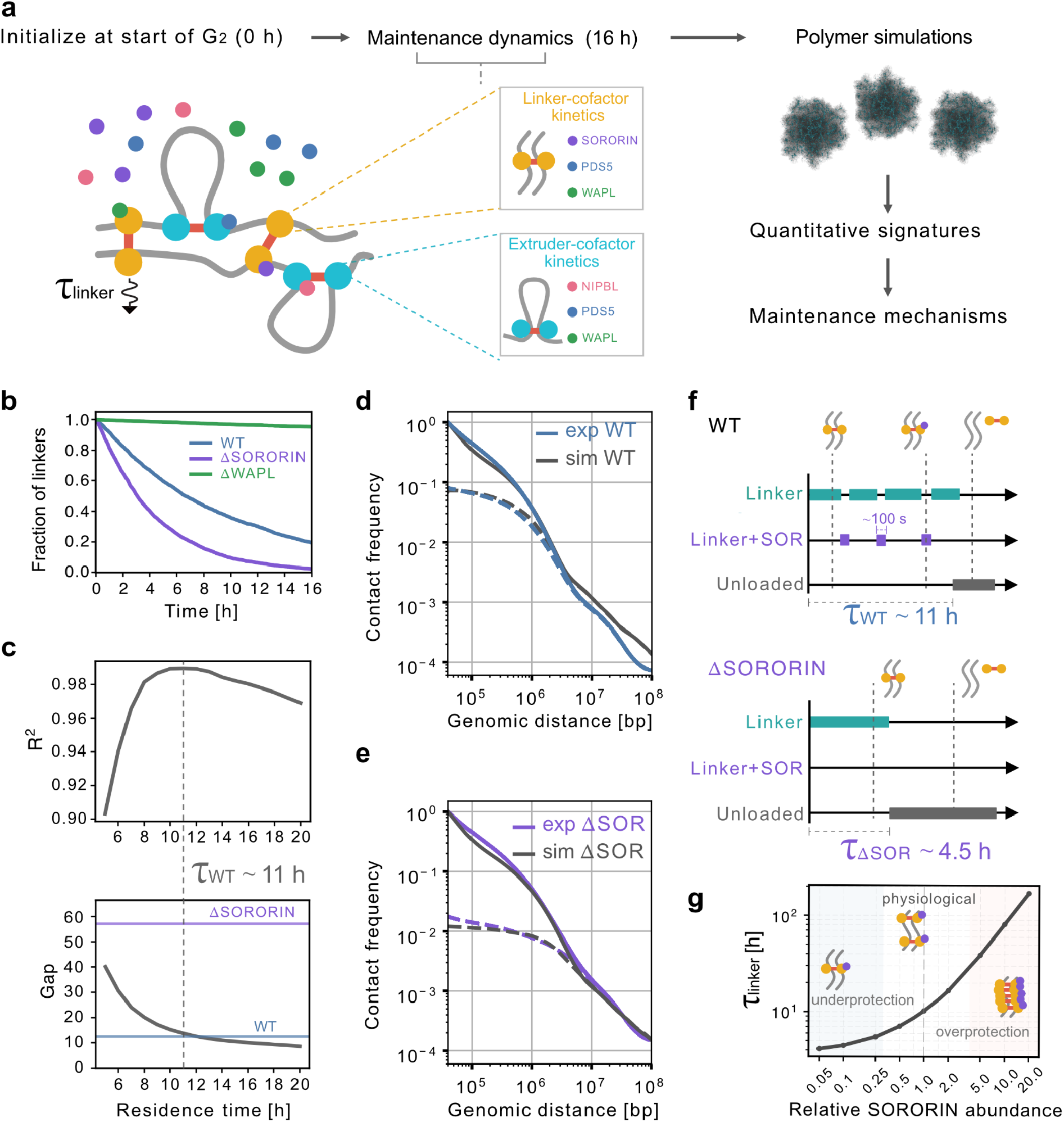
Regulatory kinetics reconcile transient cofactor binding with long-lived cohesion maintenance. **(a)** Simulation workflow. Lattice simulations were initialized from the inferred G_2_-0h (S8h) sister linker configuration (**Fig. S2**) and evolved for 16 h using chemical reaction networks for sister-linking and loop-extruding cohesins. Sister-linking cohesins transition among cofactor-bound states before unloading, whereas loop-extruding cohesins cycle between chromatin-bound and free states alongside cofactor transitions (**Fig. S3a, b**). Linker dynamics were parameterized by a residence time (*τ*_linker_). Extruder and linker positions at 16 h were then used as inputs for polymer simulations to generate P_cis_ and P_trans_ curves. **(b)** Fraction of sister-linking cohesins remaining during 16 h of simulated G_2_ maintenance in WT, ΔSORORIN, ΔWAPL cells. **(c)** WT linker residence time was inferred by comparing experimental and simulated scsHi-C. Top, combined R² for P_cis_ and P_trans_ in log-log space as a function of linker residence time. Bottom, simulated WT gap (P_cis_/P_trans_ at 40 kb); dashed horizontal lines indicate the experimental WT and ΔSORORIN gaps. Both metrics constrain the WT linker residence time to approximately 11 h (dashed vertical line). **(d–e)** Contact-frequency curves from simulations using the inferred kinetics and experimental scsHi-C (*21*) for WT **(d)** and ΔSORORIN **(e)**. The ΔSORORIN curves were predicted from the WT-inferred kinetics without further fitting. Solid and dashed lines denote P_cis_ and P_trans_, respectively. **(f)** Illustration of SORORIN protection by transient sequestration of cohesin prior to unloading. In WT (top), sister-linking cohesins enter SORORIN-associated protected states that last ∼100 s, whereas their overall residence time is approximately 11 h. Loss of SORORIN (bottom) eliminates transitions into the protected state and shortens the residence time to ∼4.5 h. **(g)** Dependence of linker residence time on relative SORORIN abundance. The inferred physiological regime lies between underprotection at low SORORIN levels and overprotection at high levels.

Cofactor residence times and occupancies were parameterized using curated FRAP data, including SORORIN’s relatively rapid turnover (∼100 s (*10*)). We then ran simulations from G_2_ entry (0 h) to the experimental timepoint when the WT scsHi-C was obtained (16 h). This reaction network predicts how unloading reduces the number of linkers over time and following perturbation to cofactor abundances (**Fig. 3b**). ΔWAPL slowed linker loss, whereas ΔSORORIN accelerated it. Chromatin-bound SORORIN decreased more slowly than the linker population, such that the fraction of remaining linkers sequestered by SORORIN increased during G_2_ maintenance (**Fig. S3g,h**).

We varied linker residence times (*τ*) over a broad range and quantified agreement between simulated and experimental P_cis_ and P_trans_ using R^2^ in log-log space (**Fig. 3c, Methods**). Agreement with WT scsHi-C was high for linker residence times of ∼9-13 h and highest around 11 h, which left ∼1 sister-linking cohesin per megabase at the experimental time point (**Fig. 3c,d**). Shorter residence times resulted in greater linker loss and gaps that were too large, whereas longer residence times retained more linkers and produced gaps that were too small (**Fig. 3c**). WT scsHi-C thus enables quantitative inference of an ultra-long timescale of cohesion maintenance.

### SORORIN depletion reveals sequestration as a mechanism of cohesion protection

We next asked how transient SORORIN binding can sustain cohesion over much longer timescales (**Fig. 3b, e**). In our chemical reaction network model, SORORIN protects chromosome-bound linkers from unloading while leaving extrusion dynamics unchanged. Starting from the WT-inferred residence time (∼11 h), the model predicts a 2.5-fold shorter linker residence time for ΔSORORIN (∼4.5 h). Without further fitting, simulations using these predicted dynamics reproduced the increased gap and slight extension of P_cis_ observed experimentally in ΔSORORIN scsHi-C (**Fig. 3e**). Shorter or longer residence times did not reproduce the ΔSORORIN gap, confirming our estimated WT linker residence time **(Fig. S4a-d**).

Together, WT and ΔSORORIN support a model in which SORORIN binding repeatedly and transiently sequesters chromosome-bound linkers from WAPL-mediated unloading. By reducing the flux through the unloading pathway, short-lived SORORIN association can prolong cohesion maintenance for multiple hours. We find that protection depends primarily on the fraction of cohesins sequestered by SORORIN rather than precise details of cofactor exchange (**Fig. S4e**). Our model further predicts that increasing SORORIN above physiological concentrations would prolong linker residence time and eventually produce overprotection that could resemble a WAPL depletion phenotype in mitotic spreads (**Fig. 3g**). Thus, in contrast to a binary switch model of SORORIN function, transient sequestration reconciles the rapid kinetics of SORORIN association with its ability to provide long-lasting protection.

### WAPL depletion exposes the consequences of cohesin encounters

To test whether additional mechanisms dictate the distribution of extrusive and linking cohesins during maintenance, we turned to ΔWAPL cells in which the unloading pathway is largely disabled (**Fig. 4a**). Disabling unloading increases cohesin residence times and leads to larger extruded loops, thereby creating a collision-rich regime for extruders. This regime produces a characteristic ‘vermicelli’ chromosome morphology, with chromatin loops emanating from an axial accumulation of cohesins (*32*). WAPL depletion after cohesion establishment, with or without prior SORORIN depletion, thus provides a setting where the consequences of extruder-linker encounters may become more apparent (depletion timeline, **Fig. S5a**). Immunofluorescence imaging revealed a single cohesin axis for ΔWAPL, whereas ΔSORORIN+ΔWAPL cells displayed two largely separated axes (*6*). In scsHi-C, ΔWAPL increases the gap and shifts the P_cis_ shoulder toward larger genomic distances, and ΔSORORIN+ΔWAPL displays similar, albeit more pronounced, changes (ref. (*6*); **Fig. 1b, c**). Simulations mirroring the WAPL depletion protocol, however, failed to reproduce two experimental features of scsHi-C: (i) the gap remained too small, and (ii) the P_cis_ shoulder did not extend sufficiently toward larger genomic distances (**Fig. 4a, Fig. S5a-c**). Based on the mechanism-agnostic simulations (**Fig. 2b,e**), these discrepancies corresponded to insufficient linker separation and extruded loop size, respectively. We therefore considered processes that could account for each discrepancy.

**Figure 4.**
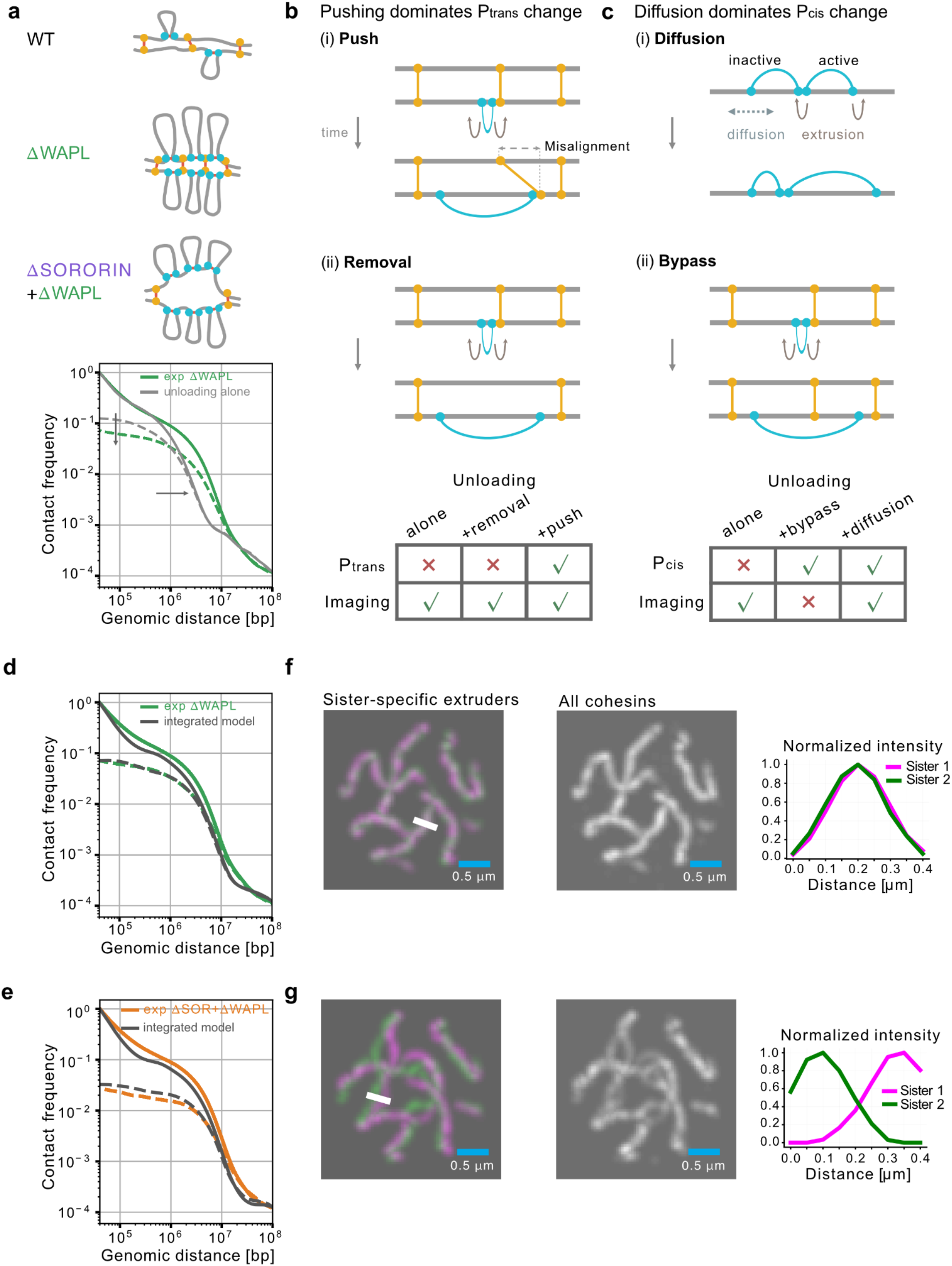
WAPL depletion resolves otherwise obscured mechanisms of chromosome organization. **(a)** *Top:* Schematic of chromosome organization in WT, ΔWAPL, and ΔSORORIN+ΔWAPL cells, illustrating the absence, formation, and separation of cohesin-rich chromosome axes, respectively. *Bottom:* Simulations with linker unloading alone do not reproduce the experimental ΔWAPL P_cis_ and P_trans_ curves, indicating that additional mechanisms must decrease the effective number of linkers (vertical arrow, increased gap) and increase extruded loop size (horizontal arrow, rightward shoulder shift). **(b)** *Top:* both pushing (i) and collision-mediated removal (ii) by extruders reduce the number of effective linkers, while pushing additionally increases linker misalignment. Curved arrows indicate extruder motion during loop extrusion. *Bottom:* Agreement between simulated mechanisms and experimental P_trans_ and imaging for ΔWAPL. Only unloading plus moderate pushing is consistent with both. **(c)** *Top:* both extruder diffusion (i) and linker bypass (ii) can increase extruded loop size and shift P_cis_ toward larger genomic distances. Bidirectional dashed arrows indicate extruder motion via diffusion. *Bottom:* Agreement between simulated mechanisms and experimental P_cis_ and imaging for ΔWAPL. Both bypass and diffusion can reproduce the P_cis_ shift, whereas bypass fails to reproduce the single-axis morphology observed by imaging. **(d–e)** Contact-frequency curves from simulations using the integrated model and experimental scsHi-C (*6*) for ΔWAPL **(d)** and ΔSORORIN+ΔWAPL **(e)**. Starting from the ΔWAPL inferred dynamics, the model predicts the ΔSORORIN+ΔWAPL curves without further fitting. Solid and dashed lines denote P_cis_ and P_trans_, respectively. **(f–g)** Simulated imaging and corresponding cohesin intensity profiles for ΔWAPL **(f)** and ΔSORORIN+ΔWAPL **(g)**. *Left:* Extruders on the two sister chromatids are shown separately in magenta and green; the white line indicates the position of the intensity line scan. *Middle:* The same simulation is shown with all cohesins (linkers and extruders) combined in gray, as would be accessible experimentally. *Right:* Normalized intensity profiles along the indicated line scan (white). The inferred dynamics reproduce largely overlapping sister axes in ΔWAPL and separated axes for ΔSORORIN+ΔWAPL.

We first asked what mechanisms could sufficiently increase linker separation to reproduce the ΔWAPL gap. We considered two major extruder-linker encounter rules proposed previously: (i) pushing of linkers by extruders (*15*, *17*), and (ii) collision-mediated removal of linkers (*16*) (**Fig. 4b**). In our simulations, both mechanisms increased the effective separation between linkers (**Fig. S6**). Sweeping both encounter rules identified a range of pushing-removal combinations that reproduced the ΔWAPL gap (**Fig. 4b**, **Fig. S5d,e**), indicating that the gap alone could not distinguish the two mechanisms. However, pushing increased linker misalignment in addition to separation. Since misalignment flattens the P_trans_ slope, it provides a potential additional constraint for distinguishing pushing from collision-mediated removal.

We next asked which mechanisms could increase extruded loop size in ΔWAPL beyond the effect of prolonged extruder residence times. We considered two candidate mechanisms: (i) extruder diffusion and (ii) linker bypass by extruders. Both mechanisms increased the mean extruded loop size and produced a rightward shift in P_cis_ (**Fig. 4c**). However, the bypass probabilities required to reproduce the ΔWAPL P_cis_ shift led to splitting of sister axes in simulations and were inconsistent with chromosome morphology observed by imaging (**Fig. 4c**; **Fig. S7**). In contrast, extruder diffusion reproduced the P_cis_ shift by increasing the loop size while preserving the observed chromosome morphology (**Fig. S8a-d**). Due to cofactor exchange, extruders cycle between active and inactive states (*28*) (**Fig. S8a**). Together with cofactor cycling, diffusion allowed collided loops held by active extruders to continue growing while those held by inactive extruders were gradually lost, generating a ‘tug-of-war’ for the loop sizes of collided extruders. Thus, although extruder diffusion had little effect on WT chromosome folding simulations (*28*), it becomes consequential in the collision-rich ΔWAPL regime.

We therefore combined candidate mechanisms to determine their relative contributions using ΔWAPL chromosome folding (**Methods**, **Fig. S9**). Jointly modeling these mechanisms narrowed the solutions to moderate pushing together with a smaller contribution from collision-mediated removal (**Fig. S9**). The resulting integrated model reproduced both P_cis_ and P_trans_ in ΔWAPL and, without additional fitting, predicted the ΔSORORIN+ΔWAPL scsHi-C profile (**Fig. 4d,e**).

To validate the integrated model, we tested whether it could also reproduce chromosome morphology observed experimentally by microscopy. In agreement with experiments, simulated imaging of cohesin positioning displayed a unified cohesin axis in ΔWAPL and split axes in ΔSORORIN+ΔWAPL (**Methods**, **Fig. 4f,g**). Importantly, perturbations to the integrated model yielded morphologies inconsistent with imaging (**Fig. S10**). For example, although reproducing the ΔWAPL scsHi-C gap required increased separation between linkers, excessively large linker separations (>200 Mb) on the ΔWAPL background produced fully resolved sister chromatid axes, instead of the single axis observed experimentally (**Fig. S9d**).

Together, these results show that cohesin unloading alone is insufficient to account for chromosome folding in WAPL mutants. In this collision-rich regime, extruder diffusion and extruder-linker encounters become important determinants of sister-chromosome conformation.

### Perturbations expose otherwise hidden cohesin activities during maintenance

Why were mechanisms inferred in ΔWAPL not evident in WT and ΔSORORIN data? Under WT extruder dynamics, pushing and diffusion at the levels inferred from ΔWAPL produced little detectable change in chromosome folding (**Fig. S11a**). Similarly, even free bypass and collision-mediated linker removal could reproduce WT chromosome folding (**Fig. S11b,c**). Thus, these processes leave little detectable imprint on chromosome folding under WT dynamics but become apparent when WAPL depletion suppresses unloading and creates a collision-rich regime.

We next asked whether the presence of additional encounter mechanisms affected the linker residence time inference under WT conditions. Allowing linker residence time and pushing resistance to vary jointly did not alter this inference: the best agreement remained at 10-12 h across a broad range of pushing resistance (**Fig. S11d-f**). Still, WT folding can rule out very strong pushing, as this generates excessive linker misalignment and overly flat P_trans_ (**Fig. S11f, g, h)**. In the resulting pushing regime, linker misalignment increased only modestly during prolonged G_2_, from ∼141 kb at entry to ∼189 kb by 16 h. The majority of linker misalignment is therefore predicted to accumulate during S-phase, consistent with the observation that the slope of P_trans_ changes substantially during S phase but remains relatively similar throughout prolonged G_2_ arrest (**Fig. S11i**).

Together, these analyses show that WT chromosome folding largely masks mechanisms beyond unloading during maintenance (**Fig. 5b**). These otherwise hidden processes are revealed by comparing simulated and experimental chromosome folding under perturbations.

**Figure 5.**
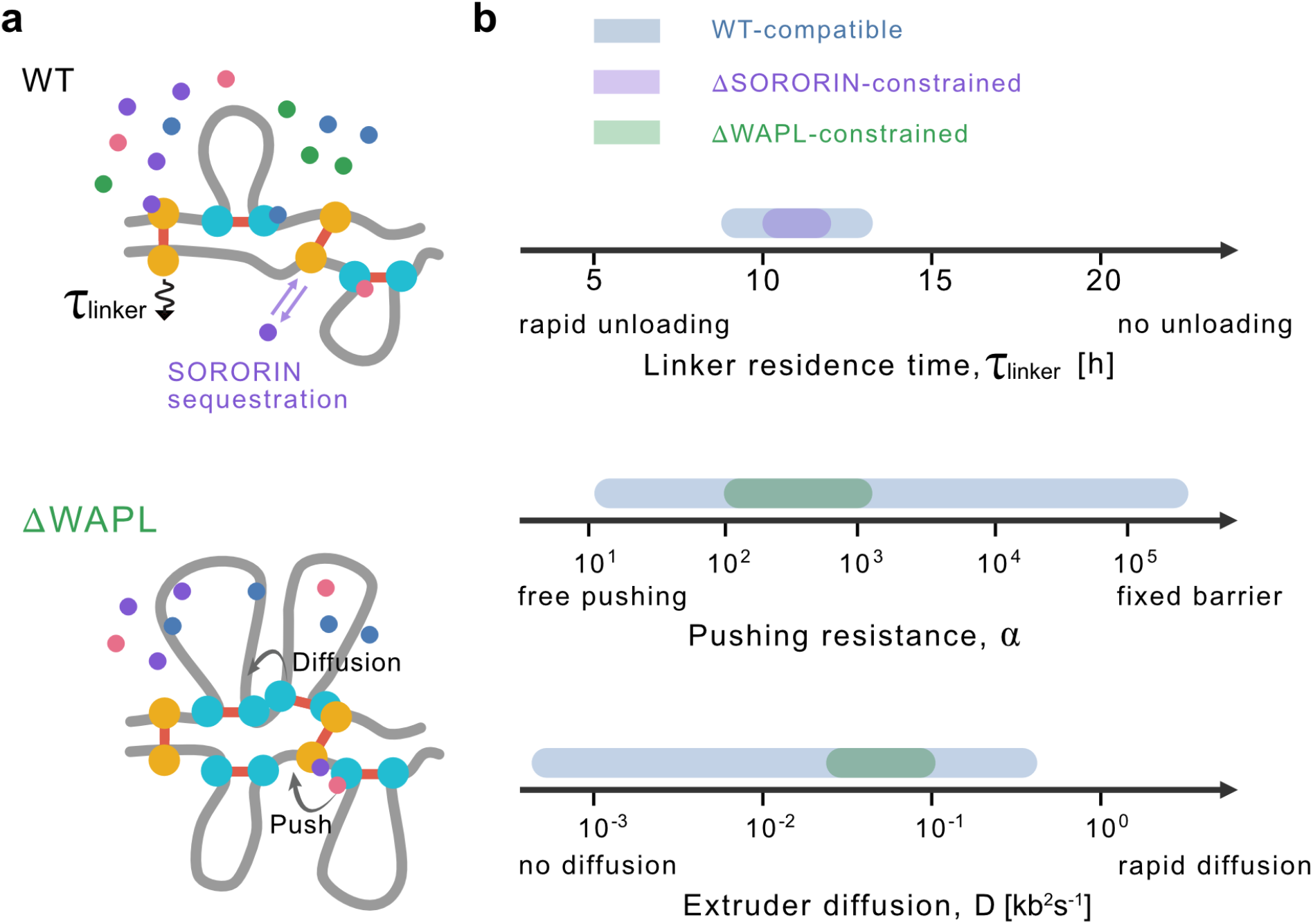
Cofactor perturbations expose otherwise hidden cohesin maintenance dynamics. **(a)** Schematic of cohesin dynamics during maintenance. SORORIN sequestration protects sister-linking cohesins from unloading, while pushing and extruder diffusion shape sister chromosome folding. **(b)** Constraints on linker residence time, pushing resistance, and extruder diffusion. WT chromosome folding weakly constrains these parameters, while ΔSORORIN and ΔWAPL provide additional constraints.

## Discussion

Our framework provides mechanistic insight into cohesin activities and their regulation, which remain intractable to direct observation. We demonstrate that 3D chromosome organization as profiled by scsHi-C contains quantitative information about cohesin linkers, extruders, and their dynamics. Specifically, while within-chromatid folding primarily reflects extrusion dynamics, between-sister folding largely reflects linker positioning. We use these complementary features to determine how cohesin unloading, diffusion, and pushing shape replicated chromosome organization. This approach uncovers new aspects of how sister-linking cohesins persist, how loop-extruding cohesins move, and how these two populations interact.

Our analysis indicates that sister-linking cohesins unload slowly but continuously throughout G_2_. We infer a residence time of ∼11 h, somewhat longer than previous estimates of 6-8 h from double-exponential fits to FRAP data (**Table S2**) (*3*, *33*). The linker reaction-network model attributes this long residence time to a lower rate of WAPL-dependent unloading than inferred for loop-extruding cohesins in G_1_ (*28*). The inferred network further predicts differential cofactor association, with PDS5 preferentially associated with sister-linking cohesins and WAPL with loop-extruding cohesins (**Fig. S3f**). This prediction is consistent with the preferential association of PDS5 with acetylated cohesin (*34–37*) and the increased resistance of acetylated cohesin to WAPL-dependent unloading (*32*, *38–40*). Intriguingly, the residence time inferred for sister-linking cohesins after SORORIN depletion (∼5 h) is also similar to the measured lifetime of stable, acetylated STAG1-containing cohesin in G_1_ (*41*). Together, these observations raise the possibility that long-lived cohesin populations in G_1_ and G_2_ share a common molecular basis involving cohesin acetylation and reduced WAPL-dependent unloading.

Our findings establish transient SORORIN sequestration as a mechanism of cohesion protection. Repeated sequestration reduces WAPL-dependent unloading of sister-linking cohesins and is sufficient to stabilize linkers without requiring a long-lived SORORIN-cohesin interaction. Although SORORIN is essential for maintaining cohesion, our results argue that it does not act as a simple lock for cohesion stability. Instead, we infer that SORORIN acts as a modest stabilizer, increasing cohesin lifetime by ∼2.5-fold at physiological concentrations. Our simulations are consistent with SORORIN sequestration operating throughout S phase, and recapitulate the observed reduction in stably bound cohesin at the start of G_2_ if SORORIN is depleted at the start of S phase (*7*). Because the predicted cohesin lifetime remains several hours after SORORIN depletion, our analysis suggests that the rapid removal of arm cohesion in prophase requires additional stimulation of cohesin unloading beyond loss of SORORIN protection alone (**Fig. S4f**), consistent with the action of multiple potential pathways (*42–44*). Dynamic SORORIN protection via sequestration therefore reconciles the long-term maintenance of cohesion with its rapid removal in prophase, avoiding the kinetic constraint imposed by a stably bound protective factor. Indeed, the concentration dependence provided by the sequestration mechanism makes cohesin lifetime tunable through changes in SORORIN abundance. Such tunability could in principle contribute to spatially regulated cohesion protection at centromeres (*45*) while still allowing cohesion to be rapidly released when sister chromatids must segregate.

By substantially extending cohesin residence times and increasing cohesin density, WAPL depletion exposes processes whose effects are negligible in wild-type settings. Moderate pushing captures changes in inter-sister contacts, whereas diffusion can explain the extension of long-range contacts within chromatids. The inferred extruder diffusion coefficient (D ≈ 0.031 kb^2^s^-1^) is lower than that measured for naked DNA *in vitro* (∼ 0.4 kb^2^s^-1^) (*46*). This reduced mobility *in vivo* is consistent with *in vitro* observations that even individual nucleosomes impede cohesin diffusion along DNA (*46*, *47*). Low levels of extruder bypass or collision-mediated removal remain difficult to resolve from the current data, leaving open whether either process occurs at low frequency. More generally, the effects of pushing, diffusion, and other encounter-dependent processes may accumulate the longer cohesins remain chromosome bound. Such cumulative effects may thus become increasingly important in physiological settings with prolonged cohesin residence, including pro-B cells (*48*), where Pax5 represses WAPL expression, and oocytes (*49*, *50*), where sister-linking cohesins can persist for years.

The level of pushing that we infer provides insight into the origin of linker misalignment. Since the inferred increase in misalignment during G_2_ is small relative to the misalignment already present at its onset, our analysis indicates that the majority of linker misalignment emerges during S phase. The mechanism generating this misalignment, whether through cohesin-fork interactions during replication (*17*), transcription (*51*), or transient structural perturbations during replication termination (*14*), will be an interesting topic for future studies.

The current approach has several limitations. While the identified network quantitatively captures both the chemical kinetics and 3D conformation across WT and perturbed cells, the underlying regulatory architecture is likely to be more complex than represented. Limited kinetic measurements restricted our analysis to minimal networks that do not explicitly model cohesin acetylation and deacetylation dynamics (*52–55*). Uncertainty in measured cofactor abundances and kinetics also propagates to the inferred reaction rates. Resolving additional regulatory network details will require precise, time-resolved measurements of cofactor abundance and kinetics before and after perturbation. The lattice model also coarse-grains processes that can influence extruder-linker encounters. For example, the effects of CTCF (*18*) and linker diffusion (*17*) are captured through the effective resistance to pushing. The model approximates established linkers as randomly positioned, whereas sister-linking cohesins can preferentially accumulate in discrete cohesion zones (*18*). Explicitly modeling these features could enable inference of their individual contributions. Extending the framework to centromeric cohesion would likely require additional regulation by Shugoshin (*45*).

In summary, the framework presented here enables quantitative inference of the molecular mechanisms by which cohesion, loop extrusion, and associated cofactors shape replicated chromosome organization. More broadly, the dominance of one mechanism in shaping chromosome folding does not imply the absence of others. Perturbations can expose otherwise hidden mechanisms of chromosome folding that leave little detectable signature under unperturbed conditions. Together, these findings provide a foundation for dissecting how cohesin-associated activities contribute to key cellular processes including DNA replication (*56*), homology search during repair (*19*, *20*), and chromosome segregation (*6*, *16*).

## Methods

### Cohesin biochemical kinetics

#### Literature-derived model parameters

Quantitative parameters for the biochemical reaction networks were obtained from measurements of cohesin abundance and chromatin-binding kinetics. Total copy numbers of RAD21 and its regulatory proteins were estimated by averaging values obtained from fluorescence correlation spectroscopy (*33*) and mass spectrometry (*57*). Live-cell imaging identified approximately equal stable chromatin-bound, dynamic chromatin-bound, and soluble RAD21 populations in G_2_ (*3*). Absolute protein abundances were scaled to the simulated chromosome size (800 Mb per sister chromatid) using the genome size (19.539 Gb) (23). This yielded an estimated mean separation of approximately 206 kb between sister-linking cohesins, consistent with the estimate in ref. (*58*). Protein abundances, chromatin-bound fractions, and residence times used to parametrize the reaction networks are summarized in **Table S1**.

Binding kinetics parameters, including chromatin-bound fractions and residence times, were obtained from FRAP measurements: WAPL, PDS5, and SORORIN from ref. (*10*); NIPBL from ref. (*59*). The residence time of the stable chromatin-bound RAD21 population was estimated to exceed 5 h (see **Table S2**) based on biexponential fits to long-term FRAP measurements (*3*, *33*) and acetylated SMC3 turnover assays (*53*). A comparable residence time has been reported for the stable population of STAG1-containing cohesin in G_1_ (*41*). Given the uncertainty in the estimate of residence time for sister-linking cohesin, this parameter was treated as a free variable in the model. In contrast, the residence time of loop-extruding cohesin was fixed at 600 s in G_2_ based on FRAP measurements (*33*).

#### Reaction network formulation

The reaction network was constructed based on the following experimentally supported assumptions, adapted from the extruder framework (*28*). Sister-linking and loop-extruding cohesins were modeled as distinct regulatory populations, motivated by their differential cofactor associations and susceptibility to WAPL-dependent release (*2*, *18*, *21*, *40*, *58*).

1. Sister chromatid cohesion is maintained by monomeric cohesin complexes (*58*, *60*).
2. The core cohesin complex, consisting of SMC1, SMC3, RAD21, and either SA1 (STAG1) or SA2 (STAG2), is modeled as a single species that associates with and dissociates from chromatin as a complex (*1*, *61*).
3. NIPBL associates predominantly with loop-extruding cohesins and negligibly with sister-linking cohesins (*62*).
4. Stable chromatin association of SORORIN, PDS5, and WAPL requires the core cohesin complex.

● SORORIN chromatin association is cohesin dependent (*7*), consistent with extensive overlap between SORORIN ChIP-seq peaks and SMC3-binding sites (*5*, *10*).
● PDS5 chromatin association is cohesin dependent (*63–65*).
● WAPL chromatin association is cohesin dependent (*66*).
5. The stoichiometric coefficient of cofactors to sister-linking cohesin is set to 1.

● SORORIN: A 1:1 stoichiometry with sister-linking cohesin has been demonstrated by single-molecule imaging (*58*).
● PDS5 and WAPL: Direct single-molecule measurements of stoichiometry are unavailable. However, proteomic measurements indicate that both cohesin cofactors are sub-stoichiometric relative to RAD21, arguing against the need to include oligomeric cofactor association in the model (*67*, *68*).
6. Association of SORORIN and WAPL with cohesin is PDS5 dependent (*5*, *8*).

#### Parameterization of WT reaction kinetics

The interactions between cohesin and its regulators were modeled as a chemical reaction network governed by mass-action kinetics (**Fig. 3a, Fig. S3a**). A system of coupled ordinary differential equations describing the concentrations of extruders, linkers, PDS5, WAPL, SORORIN, and NIPBL was derived (**Supplementary Note 1**). Symbols used in the reaction-network formulation are summarized in **Table S3**.

FRAP measurements for PDS5 and WAPL report the combined chromatin-bound fractions of molecules associated with both loop-extruding and sister-linking cohesins (**Table S1**). To separate these two populations, we allowed the chromatin-bound fractions of PDS5 and WAPL to differ between extruders and linkers while constraining their sum to equal the experimentally measured total bound fraction.

Specifically, PDS5 and WAPL were allowed to have different association rates (*k*_on_) with extruders and linkers but the same dissociation rate (*k*_off_). Thus, the bound fractions for PDS5 at linkers 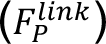 and extruders 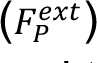 could differ, subject to the constraint 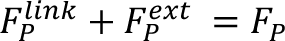, where *F_P_* is the experimentally measured total bound fraction. The same constraint was applied to WAPL. Together with the requirement that all reaction rates be non-negative, these constraints defined the feasible region for partitioning PDS5 and WAPL between sister-linking and loop-extruding cohesins (**Fig. S3e**). The centroid of the feasible region (0.120, 0.302) was used to parametrize transition rates for the WT linker and extruder networks. The resulting reaction rates are summarized in **Table S4**. Full derivations of the kinetic equations are provided in **Supplementary Note 1**.

##### Determination of reaction kinetics for depletion conditions

After parametrizing the WT reaction network, the temporal evolution of sister-linking cohesins and associated cofactors was obtained by numerical integration of the ODE system. Perturbation conditions (ΔSORORIN, ΔWAPL, and ΔSORORIN+ΔWAPL) were simulated by modifying the abundance of the depleted regulator according to the corresponding experimental depletion protocols (**Fig. S2a** and **Fig. S5a**) (*6*, *21*). Each perturbation condition was simulated over the corresponding experimental time course. Transition probabilities extracted at the midpoint of each experimental interval were subsequently used as input for the multistate extrusion model described previously (*28*). Depletion efficiencies of 99% for WAPL and 97% for SORORIN were used as reported previously (*6*).

##### Determination of the initial density of linkers for ΔSORORIN simulations

Because scsHi-C measurements immediately after DNA replication (G_2_-0h) were unavailable for SORORIN depletion, we estimated the initial density of sister linkers using a minimal birth-death model describing linker establishment during S phase. The model assumes that linkers are established at a constant rate *λ* and unloaded with condition-specific residence times *τ*_W*T*_ and *τ_Δs_* for WT and ΔSORORIN cells, respectively. Starting from *N*(0) = 0 at S-phase entry, the linker density evolves as 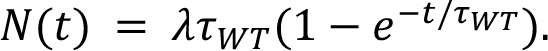. The establishment rate *λ* was determined from the WT linker density inferred by fitting the G_2_-0h scsHi-C data, yielding *λ*= 0.64 h^-1^Mb^-1^. Assuming that SORORIN depletion alters only the residence time while leaving the establishment rate unchanged, the same model was used to estimate the initial linker density (∼2.4 Mb^-1^) for ΔSORORIN simulations. This estimate is also consistent with the reduced abundance of stably chromatin-bound cohesin measured by FRAP in G_2_ following SORORIN depletion initiated before S phase progression (*7*).

### 1D Lattice model of linker and extruder dynamics

#### Initialization of sister linkers

Sister linkers were initialized on a pair of one-dimensional lattices representing the two sister chromatids, with positions determined by the specified spacing and misalignment distributions.

To generate non-uniform linker positions, gap sizes were sampled from either a uniform distribution or a truncated power-law (Levy-like) distribution, *p*(*g*) ∼ *g*^−£^, with *g* in [1, 10000] and *ξ* = 1.5. Truncation preserved the long-tailed spacing distribution while maintaining a fixed number of sister linkers within the simulated lattice (**Fig. 2c**).

To generate linker misalignment, sister linker pairs were first positioned independently on the two chromatids. For each sister linker, a random variable [-1, 1] was drawn to determine which anchor to displace, and the selected anchor was shifted according to the specified misalignment distribution. Because this procedure could introduce crossings, we subsequently reordered the indices and reassigned pairings to preserve the genomic ordering of anchors along the two sister chromatids. Crossings were defined as pairs of sister linkers *i* and *j* such that (*i*_1_ − *j*_1_) (*i*_2_ − *j*_2_) < 0. The configuration shown in **Fig. 2d** had a linker misalignment with a standard deviation of 229 kb.

#### Rules of engagement between linkers and extruders

Three rules were implemented at each lattice step when an extruder encountered a linker. These rules and their limiting regimes are summarized in **Table S5**.

##### Pushing

The sister linker was displaced to the nearest available lattice site with probability 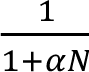, where *α* denotes the resistance to pushing and N is the number of linkers occupying the lattice site. The fixed-barrier regime was approximated by setting *α* = 50,000.

##### Collision-mediated removal

The sister linker was unloaded with probability *β* when encountered by an active extruder.

##### Bypass

An extruder bypassed the sister linker with probability *γ.* In the free-bypass regime, *γ* = 1.

#### Reaction-regulated lattice dynamics

Loop-extruding and sister-linking cohesins were simulated as a discrete-time stochastic process on a one-dimensional lattice spanning 800 Mb. WT and ΔSORORIN simulations used 320,000 lattice sites at 2.5 kb resolution with a timestep *τ*_1*D*_ =1.25 s. For ΔWAPL and ΔSORORIN+ΔWAPL simulations, a fivefold finer lattice consisting of 1,600,000 sites at 0.5 kb resolution was used, with a corresponding timestep *τ*_1*D*_ = 0.25 s. Encounter probabilities were rescaled to preserve the corresponding rates across lattice resolutions. The finer lattice resolution was used under WAPL-depleted conditions to resolve the increased frequency of cohesin-cohesin encounters resulting from higher cohesin density and increased loop processivity. Lattice dynamics were coupled to the biochemical reaction network, following the extrusion framework described previously (*28*).

At each timestep, loop-extruding cohesins could stochastically load, unload, or transition between states (*R^ext^*,*R^ext^*N, *R^ext^*P, *R^ext^*PW, *R_free_*) according to the transition probabilities obtained from the reaction network. Linkers were assigned an unloading probability determined by the corresponding condition (WT, ΔWAPL, ΔSORORIN, or ΔSORORIN+ΔWAPL). The residence time and processivity of loop-extruding cohesins therefore emerged directly from the underlying biochemical kinetics. The mean loop extrusion velocity was fixed at 1 kbs^-1^ (*28*).

### Simulation of prolonged G_2_ dynamics

To reproduce the experimental perturbation protocols, lattice simulations were performed over the corresponding G_2_ time courses for WT and each perturbation condition. The experimental conditions, perturbation timing, and scsHi-C datasets used for each condition are summarized in **Table S6**.

#### Initial conditions

WT and ΔWAPL simulations were initialized using the linker separation (∼275 kb) and misalignment (∼141 kb) inferred from scsHi-C data at G_2_-0h. Because SORORIN depletion occurs before DNA replication, ΔSORORIN and ΔSORORIN+ΔWAPL simulations were initialized using reduced linker densities estimated from the kinetic model of sister-linker establishment. Initial linker positions were obtained by subsampling the WT configuration to match the predicted linker density.

#### Simulation protocol

WT and ΔSORORIN simulations were performed using their respective biochemical parameters throughout the 16 h G_2_ simulation. To reproduce the experimental depletion protocol, ΔWAPL simulations were evolved under WT parameters for the first 6 h before switching to ΔWAPL parameters for the remaining 10 h. Similarly, ΔSORORIN+ΔWAPL simulations were evolved under ΔSORORIN parameters for the first 6 h before switching to combined depletion parameters for the remaining 10 h. The resulting lattice configurations were subsequently used as inputs for the polymer simulations described below.

#### Lattice simulations of individual encounter mechanisms

To explore each mechanism, linkers were initialized randomly without misalignment, with a mean separation corresponding to the experimentally inferred G_2_-0h configuration. Extruders were initialized at random positions independently on both sister chromatids. Simulations were first run for 2 h to equilibrate extruded loops, with linkers acting as fixed barriers to extruders (no pushing, bypass, or collision-mediated removal).

Starting from the equilibrated configurations, simulations were propagated for 16 h with each mechanism enabled individually. For unloading, linker dynamics were governed by the chemical reaction network for WAPL, PDS5, and SORORIN described above, with a basal linker residence time of 10 h and cofactor depletion set to 95%. Pushing was tested with resistance *α* = 10, 100, or 1000; collision-mediated removal with probability *β* = 0, 10^-5^, or 10^-4^; or bypass with probability *γ* = 0, 0.1, or 1. For each simulation, all other encounter mechanisms were disabled.

We then profiled how the effective number of linkers, linker misalignment, and the extruded loop size changed over time (**Fig. S6**).

### 3D Polymer model of sister chromatids

#### Polymer construction and lattice coupling

Sister chromatids were represented as two 800-Mb polymer chains, each consisting of 320,000 monomers at 2.5 kb resolution, with a bead size of *b* = 50 nm. Initial polymer conformations were generated using the grow-cubic algorithm implemented in polykit.generators.initial_conformation (https://github.com/open2c/polykit). Replication was modeled by generating a second copy of the polymer chain superimposed onto the original chain.

Extruders and linkers generated by the 1D lattice simulations were incorporated as dynamic polymer crosslinks. Sister linkers were represented as bonds connecting their corresponding loci on the two sister chromatids, whereas extruders were represented as bonds connecting monomers occupied by the two legs. Condition-specific linker and extruder positions were obtained directly from the 1D lattice simulations described above. Polymer simulations were then used to sample 3D chromosome conformations for the final lattice-derived cohesin configuration of each perturbation condition.

#### Polymer equilibration

Molecular dynamics simulations were performed using polychrom-hoomd (https://github.com/open2c/polychrom-hoomd), following the polymer simulation framework described previously (*28*). The force field included repulsive, bonded, and angular interactions, and dynamics were propagated using a Langevin integrator. WT systems were equilibrated for 5×10^6^ steps, whereas ΔSORORIN, ΔWAPL, and ΔSORORIN+ΔWAPL systems were equilibrated for 3×10^7^ steps to accommodate the slow relaxation at long genomic distances. Equilibration was followed by 10^6^ production steps for all conditions, with polymer configurations recorded every 25,000 steps to yield 40 conformations per trajectory. Five independent simulations were performed for each parameter set.

To reduce the computational cost of equilibration, WT sweep simulations were initialized from a pre-relaxed sister-chromatid backbone generated with a mean linker separation of 200 kb. The backbone was equilibrated for 10^7^ steps, followed by 5×10^6^ steps during which 200 conformations were recorded at intervals of 25,000 steps. The pre-relaxed linker bonds were removed and replaced with the linker and extruder bonds obtained from the corresponding lattice simulations. Following bond reassignment, the polymer was relaxed using the Fast Inertial Relaxation Engine (FIRE) for 5×10^5^ steps before molecular dynamics simulations.

### Parameter inference

Model parameters were inferred sequentially by comparing simulations with complementary experimental observables that constrain distinct aspects of the model. The experimental constraints used to distinguish candidate maintenance mechanisms are summarized in **Table S7**.

#### Initial sister-linker configuration (G_2_-0h)

To determine the initial sister-linker configuration at G_2_-0h, we systematically varied the linker separation (200-400 kb, 25 kb increments) and linker misalignment scale (50-175 kb, 25 kb increments). For each parameter combination, lattice and polymer simulations were performed and the resulting P_cis_ and P_trans_ were compared with the experimental scsHi-C data. The initial configuration was selected based on the combined R^2^ of the cis and trans P(s) curves over genomic distances from 40 kb to 5 Mb.

#### WT sister-linking cohesin kinetics

To determine the residence time of sister-linking cohesins during G_2_, we systematically varied the residence time from 5 to 20 h, with 5000 h included as an effectively permanent-linker limit, under minimal pushing (*α* = 50,000). Simulations were initialized from the inferred G_2_-0h linker configuration and evolved using the reaction-regulated lattice model described above. For each residence time, lattice configurations were converted into polymer conformations and compared with experimental scsHi-C data. The optimal residence time, *τ* = 11 h, was determined by maximizing the combined cis and trans R^2^ over genomic distances from 40 kb to 5 Mb.

#### Linker–extruder encounter kinetics

To determine the contributions of pushing and removal in ΔWAPL, we systematically varied the pushing resistance (*α* = 0, 5, 10, 15, 20, 50, 100, 200, 500, 1000 and 50,000) together with the collision-mediated removal probability (*β* = 10^-6^ - 10^-4^). Simulations were initialized using the residence time inferred above and evolved using the reaction-regulated lattice model. Parameter combinations were constrained by requiring the resulting linker separation, linker misalignment, and extruded loop size to be jointly consistent with experimental folding signatures. These constraints defined a feasible region in pushing and removal parameter space. The centroid of this feasible region was used to compute P_trans_.

#### Extruder diffusion

To infer the effective extruder diffusion coefficient, simulations were performed using the residence time and encounter parameters inferred above. The diffusion coefficient D was systematically varied from 0 to 0.0625 kb^2^s^-1^. For each diffusion coefficient, lattice and polymer simulations were performed and compared with experimental scsHi-C data. The optimal diffusion coefficient, D = 0.03125 kb^2^s^-1^, was determined by maximizing the cis R^2^ over genomic distances from 40 kb to 10 Mb.

#### Joint parameter optimization

Using the parameter regimes inferred above, we performed a coarse-grained joint optimization of pushing resistance (*α*), collision-mediated removal probability (*β*), and bypass probability (*γ*). To limit the computational cost, we evaluated a reduced parameter grid with *α* = 10, 100, and 1000, *β* = 10^-7^, 10^-6^, and 10^-5^, and *γ* = 10^-1^, 10^-3^, and 10^-5^. For each parameter combination, lattice and polymer simulations were performed using the diffusion coefficient inferred. The final encounter-parameter set (*α* = 100, *β* = 10^-6^, and *γ* = 10^-3^) was selected by maximizing the trans R^2^ over genomic distances from 40 kb to 10 Mb.

### Analysis of chromosome folding

#### Experimental scsHi-C analysis

Experimental scsHi-C data for HeLa cells were obtained from published datasets: WT and ΔSORORIN from ref. (*21*), and ΔWAPL and ΔSORORIN+ΔWAPL from ref. (*6*) (**Table S6**). For all conditions, contact maps from all available replicates were merged before calculating P(s). Contact probability as a function of genomic distance, P(s), was computed separately for cis and trans using the expected_cis function in cooltools (https://github.com/open2c/cooltools, ref. (*69*)). The P(s) curves were averaged across chromosomes and smoothed. All curves were normalized to the cis value at 40 kb.

#### Contact probability and P(s) from polymer simulations

To compute contact maps from polymer simulations, we processed GSD trajectory files containing 3D coordinates of two sister chromatids. A custom function was used to calculate the cis and trans contact matrices. Cis and trans contact maps were obtained by summing contacts within the two sister chromatids and between the two sister chromatids, respectively. Contact maps were computed at the single monomer level using a cutoff distance *R_c_*= 2.3b ∼ 115 nm, consistent with typical Hi-C experimental parameters (*70*). The cis and trans matrices were converted into Cooler objects using contact_maps.coolify. The average contact probability as a function of genomic distance was computed using the expected_cis function of the cooltools library.

#### Definition of key statistics

##### Gap

The gap between P_cis_ and P_trans_ is quantified as the ratio P_cis_/P_trans_ at 40 kb.

##### Trans slope

The trans slope at 40 kb is defined as the local derivative of log P_trans_(s) with respect to log s, evaluated at s = 40 kb. To reduce sensitivity to noise, P_trans_(s) is first smoothed with a Gaussian filter before taking the finite difference.

##### Cis shoulder

The cis shoulder is defined as the genomic distance at which the log-log derivative of P_cis_(s) is maximal.

##### Mean loop size

The cis mean loop size is defined as the mean genomic distance between the two legs of loop-extruding cohesins.

##### Goodness of fit

The goodness of fit is quantified by R^2^ in log-log space:

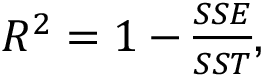

where 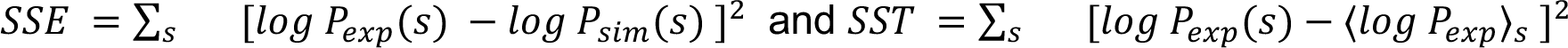. Here, *P_exp_*(*s*) and *P_sim_*(*s*) denote the experimental and simulated contact frequencies at genomic separation *s*, respectively, and ⟨*log P_exp_*⟩*_s_* denotes the mean across *s*. The mean and sums are evaluated over the genomic ranges of 40 kb-5 Mb for WT and 40 kb-10 Mb for ΔWAPL. Cis and trans values are concatenated to calculate the combined R^2^.

### Simulated imaging from polymer conformations

We followed the experimental imaging processing protocol described previously (*6*) and converted 3D polymer simulation coordinates into voxel-based fluorescence-like images. 3D simulation coordinates were discretized into cubic voxels of dimension 100 nm×100 nm×100 nm. Monomer and cohesin positions were rasterized onto a regular 3D grid using a custom voxelization procedure based on spatial binning (*28*). The resulting voxelized densities were subsequently convolved with a Gaussian filter with a width of 150 nm to approximate the diffraction-limited point-spread function of the microscope. For visualization, a 20-slice slab of the rasterized volume was selected along the imaging axis prior to projection. Individual cohesin populations on each sister chromatid were rasterized separately and converted into 2D fluorescence-like images by maximum-intensity projection along the imaging axis. Projected voxel intensities were normalized and mapped onto RGB color channels using custom green and magenta colormaps, after which the channels were overlaid to generate composite fluorescence images.

## Data and Code Availability

The scsHi-C datasets used in this study and their GEO accessions are listed in Table S6. Code supporting the analyses is publicly available at https://github.com/Fudenberg-Research-Group/sister-cohesin-dynamics.

## Acknowledgements

X.Y., M.M.C.T. and G.F. were supported by NIH Research Project Grant R35GM143116 to G.F. A.G. was supported by the European Research Council (ERC), European Union’s Horizon 2020 research and innovation programme grant agreement no. 101163751. The authors thank Fena Ochs, Elphège Nora, Daniel Gerlich, Zsuzsanna Takacs, Dmitry Mylarshchikov, Flavia Corsi, and Paul Batty for helpful discussions and feedback on the manuscript.

## Author Contributions

X.Y., A.G., and G.F. conceived of the project. X.Y. and G.F. designed the mechanistic inference approach. X.Y. developed models, implemented simulations, and performed analyses. M.M.C.T. contributed to model development and advised on simulation implementation. X.Y. drafted the initial manuscript with G.F. All authors reviewed and edited the manuscript.

## Competing Interests

The authors declare no competing interests.

## Supplementary Figures

**Figure S1.**
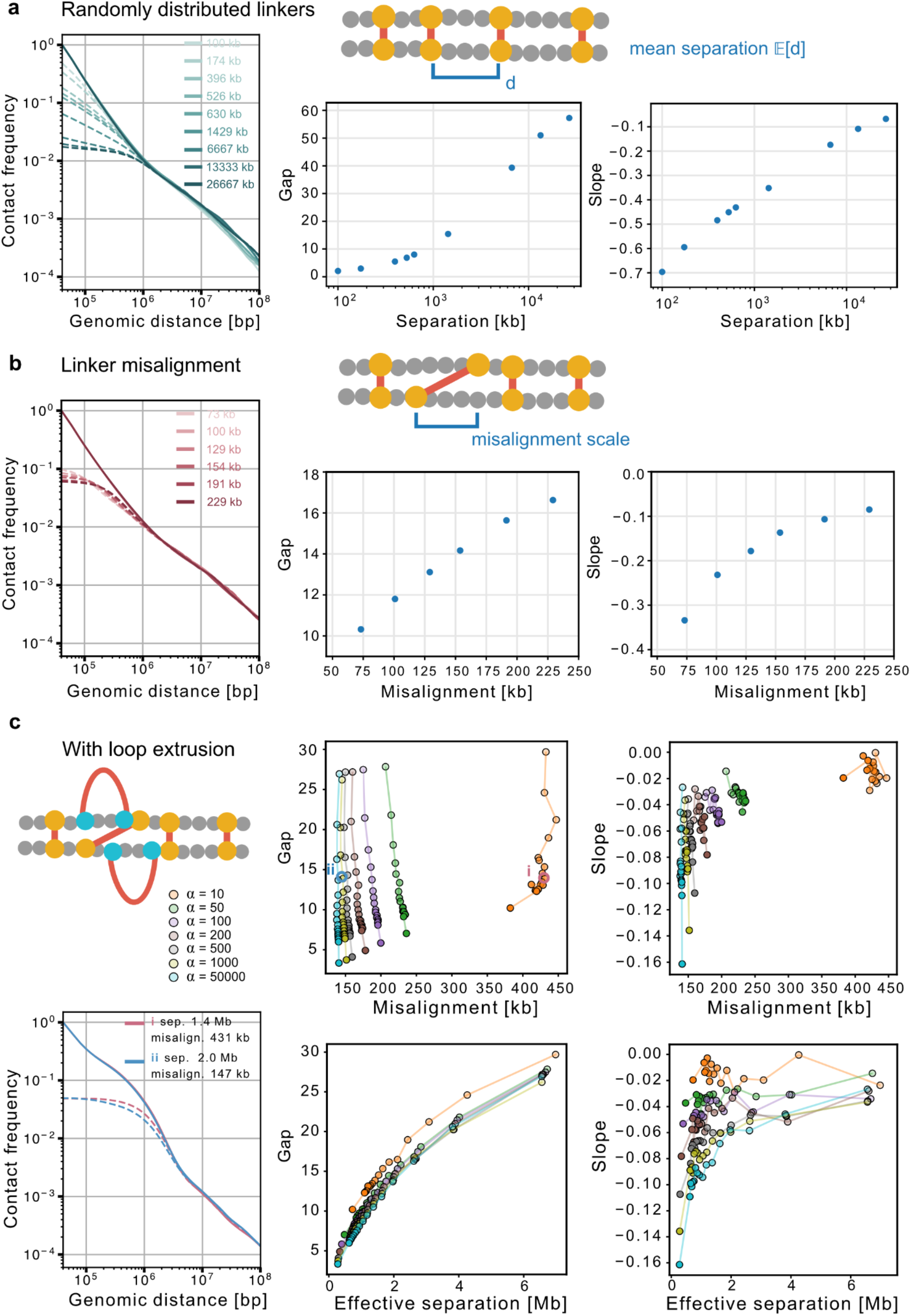
Distinct contact-frequency signatures provide complementary constraints on sister-chromatid organization. **(a)** Effect of linker separation for randomly distributed linkers. Linker positions were sampled randomly, with *d* denoting the separation between neighboring linker anchors and *E[d]* its mean. *Left:* Contact frequency versus genomic distance for a range of mean linker separations. *Right:* the corresponding gap and P_trans_ slope as a function of linker separation. **(b)** Sister-linker misalignment was introduced in simulations by sampling genomic displacements to add to paired linker positions, with the misalignment parameter controlling the width of the displacement distribution. Simulations were performed at a fixed mean linker separation of 1 Mb. *Left:* contact frequency versus genomic distance for a range of misalignments. *Right:* Corresponding gap and P_trans_ slope as a function of the degree of misalignment. **(c)** Dependence of simulated scsHi-C signatures on linker configurations with extruders present. In these simulations, the parameter *α* controls the misalignment scale. Within each *α*, shading and connecting lines indicate increasing residence time *τ* that, in combination with *α*, controls the effective linker separation. *Top:* Gap (left) and P_trans_ slope (right) plotted against linker misalignment. *Bottom:* The same quantities plotted against effective linker separation. Highlighted points (**i, ii**) indicate two configurations with similar gaps but distinct linker separations, misalignments, and P_trans_ slopes; their corresponding contact-frequency curves are shown on the lower left. Linker separation is the primary determinant of the gap, with linker misalignment providing additional modulation, whereas the P_trans_ slope provides complementary information about linker misalignment.

**Figure S2.**
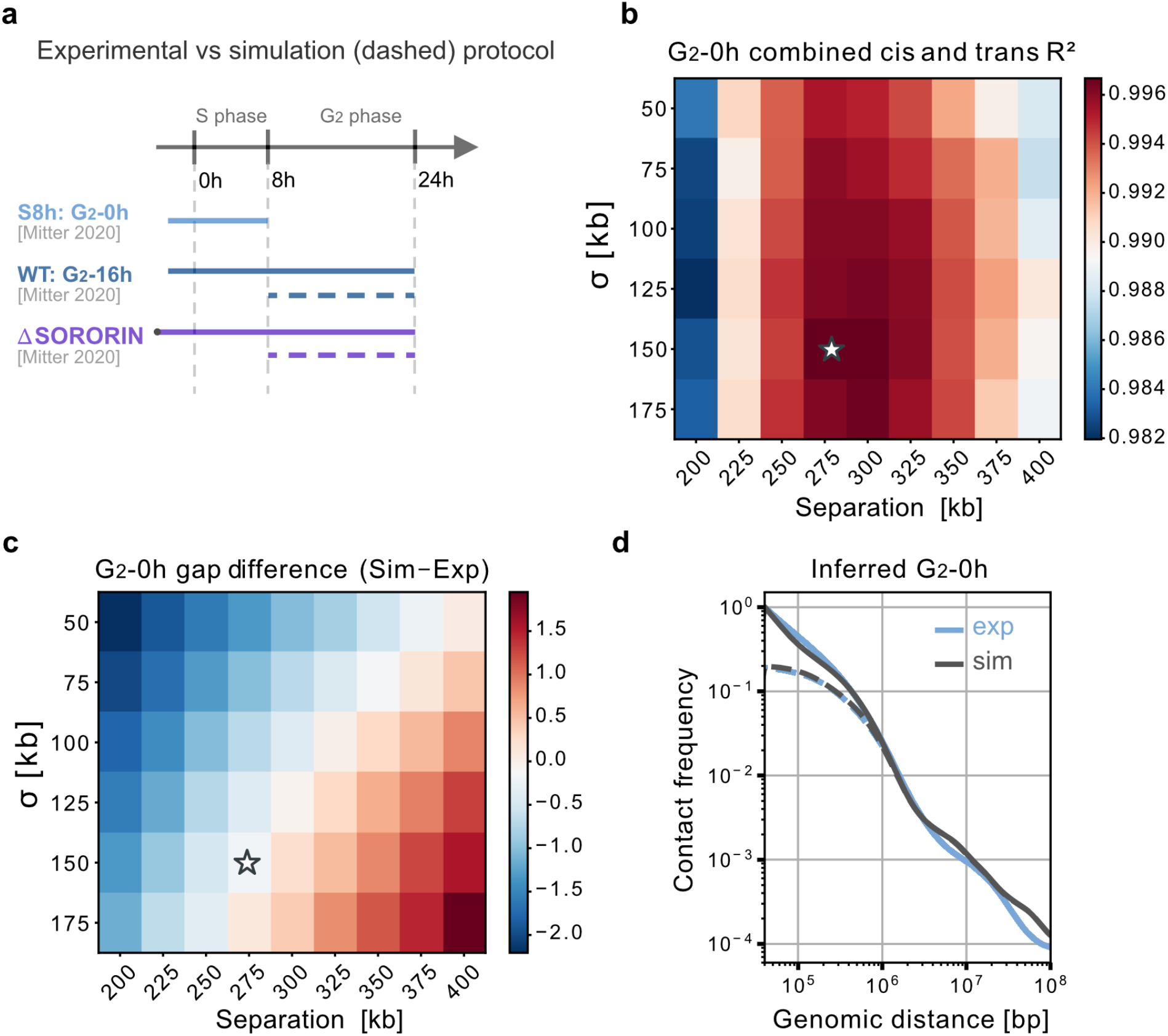
Linker configuration at G_2_-0h is inferred from scsHi-C. **(a)** Experimental protocols for time-resolved scsHi-C, reproduced from ref. (*21*). WT scsHi-C data were collected at 8 h (S8h, G_2_-0h) and 24 h (S24h, G_2_-16h) after S phase release. Simulation protocols were designed to mirror the experimental time courses. WT simulations were initialized from the inferred G_2_-0h configuration and propagated to G_2_-16h. For ΔSORORIN, SORORIN was depleted before S phase entry. Simulations were initialized with the ΔSORORIN linker abundance predicted at G_2_-0h and propagated for 16 h. **(b–c)** Inference of the G_2_-0h sister-linker configuration by jointly varying linker separation and misalignment scale, *σ*. **(b)** Combined R^2^ between experimental and simulated P_cis_ and P_trans_. **(c)** Difference between the simulated and experimental P_cis_/P_trans_ gap. Stars indicate configuration with the highest combined R^2^. **(d)** Experimental and simulated P_cis_ and P_trans_ for the best-fitting G_2_-0h sister-linker configuration. WT G_2_-0h scsHi-C were reanalyzed from ref. (*21*).

**Figure S3.**
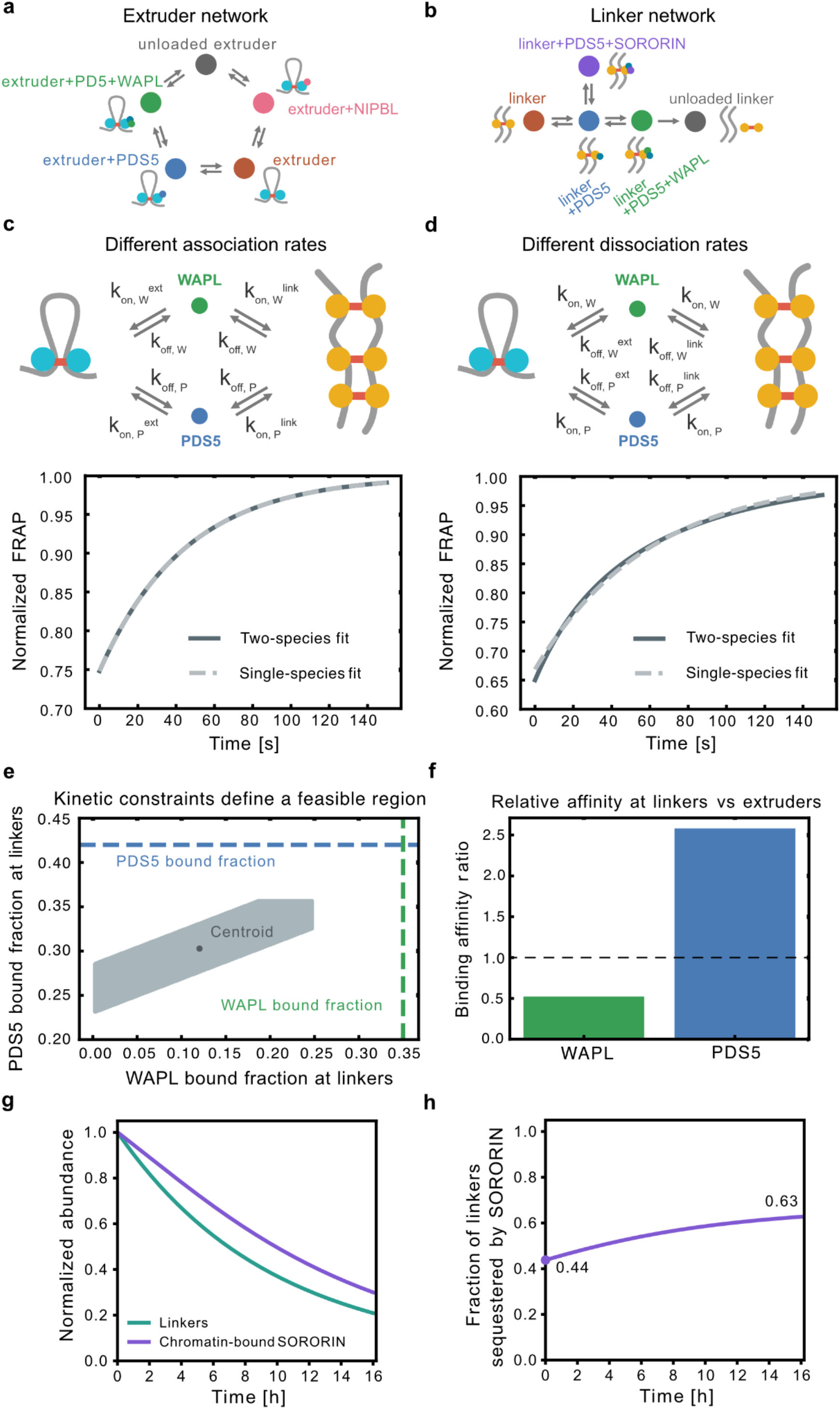
Joint parameterization of cohesin linker and extruder networks. **(a)** Reaction network describing cofactor-dependent transitions of loop-extruding cohesins. Colored circles indicate states and directional arrows indicate possible state transitions. Extruders transition in a chemical reaction cycle from free extruders to chromatin-bound states associated with NIPBL, no additional factors, PDS5, and WAPL plus PDS5 , before unloading back to the free state. **(b)** Reaction network describing cofactor-dependent transitions of sister-linking cohesins. At the start of G_2_ linking cohesins are initialized in chromatin bound states and reversibly transition between chromatin-bound states associated with SORORIN, PDS5 and WAPL before unloading into the free linkers. Since we assume linker cohesins are only specified concomitantly with DNA replication, linkers are not loaded back onto chromatin from the unloaded population (in contrast with extruders). **(c)** FRAP recovery for models with distinct association rates, *k*_on_, but a shared dissociation rate, *k*_off_, between linkers and extruders. The FRAP recovery from a single-species model (*k*_on_ = 0.00798, *τ* = 45.0 s) is reproduced by a two-species model with *k*_on_^ext^ = 0.00598 and *k*_on_^link^ = 0.00150. **(d)** FRAP recovery for models with a shared association rate, *k*_on_, but distinct dissociation rates, *k*_off_, between linkers and extruders. The FRAP recovery from a single-species model (*τ*^fit^ = 60.0 s) is reproduced by a two-species model with *τ*^ext^ = 20.0 s and *τ*^link^ = 70.0 s. **(e)** How kinetics for the combined network are identified as the center of the feasible region for the varying *k*_on_ model. Dashed blue and green lines indicate the maximum PDS5 and WAPL bound fractions, respectively. **(f)** Predicted linker-to-extruder binding-affinity ratios for WAPL and PDS5 using the parameterization selected in **(e)**. The dashed line indicates an equal affinity ratio between linkers and extruders. **(g-h)** Predicted SORORIN occupancy during G_2_ maintenance. **(g)** Linker and chromatin-bound SORORIN abundances normalized to their values at G_2_ onset. Chromatin-bound SORORIN decreases more slowly than the total bound linker population. **(h)** Fraction of remaining linkers sequestered by SORORIN. The experimentally constrained initial state corresponds to a SORORIN-sequestered fraction of 0.44, which increases to 0.63 over 16 h of simulated G_2_ maintenance.

**Figure S4.**
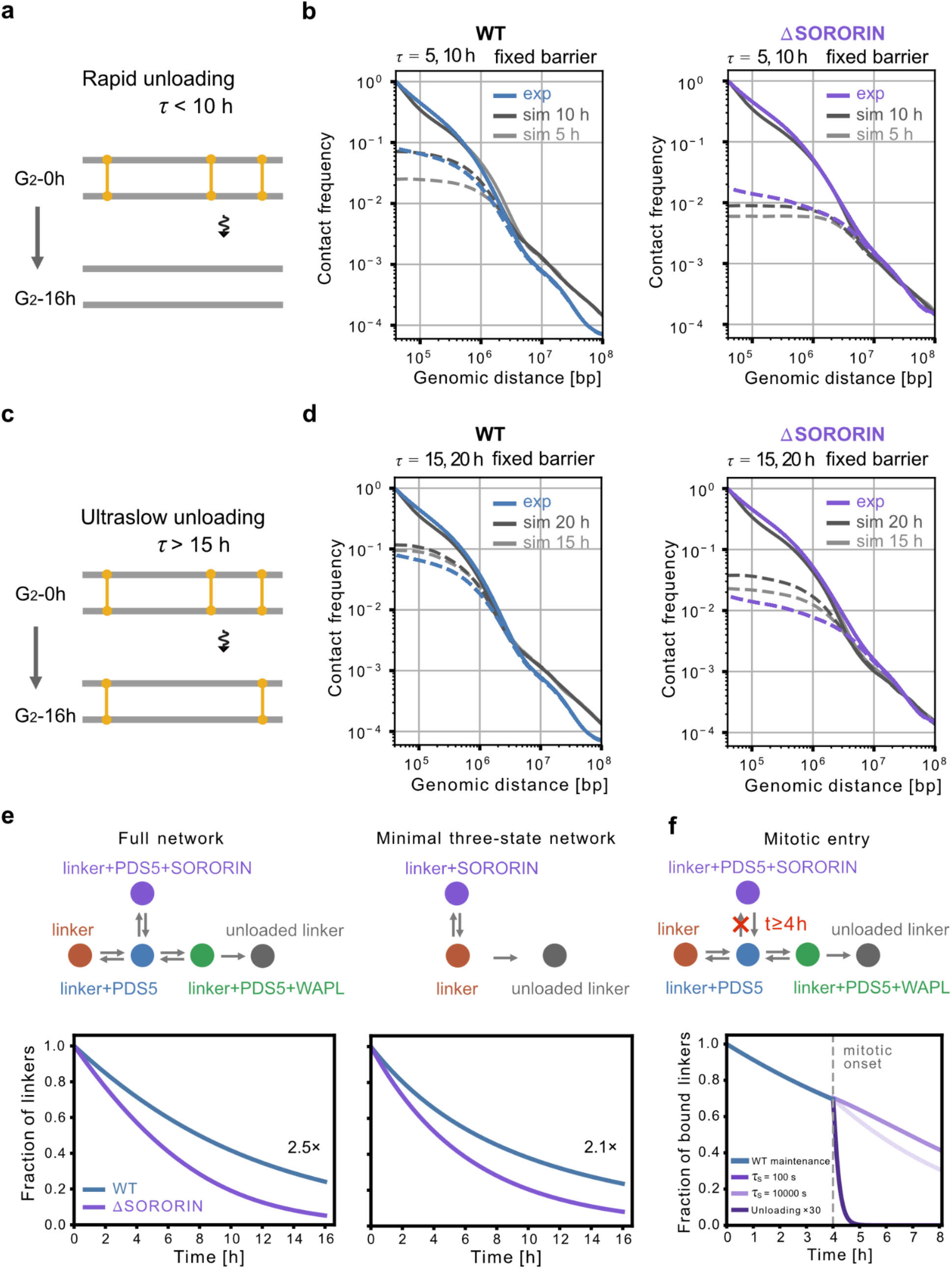
WT and ΔSORORIN scsHi-C constrain the lower and upper bounds on sister-linker residence time and support a sequestration mechanism. **(a)** Schematic of rapid linker unloading (*τ* < 10 h), which leads to extensive loss of sister-linkers during G_2_ maintenance. **(b)** WT and ΔSORORIN scsHi-C constrain the lower bound on linker residence time. With linkers acting as fixed barriers (*α* = 50,000), rapid unloading (*τ* = 5 h) produces gaps larger than observed experimentally in both WT and ΔSORORIN. Increasing the residence time to 10 h improves agreement with WT, but the ΔSORORIN gap remains overestimated. **(c)** Schematic of ultraslow linker unloading (*τ* > 15 h), which retains a larger fraction of sister linkers throughout G_2_. **(d)** WT and ΔSORORIN scsHi-C constrain the upper bound on linker residence time. Ultraslow unloading (*τ* = 15 and 20 h) retains too many sister linkers and fails to reproduce the WT scsHi-C gap. The ΔSORORIN gap is likewise underestimated. Together, WT and ΔSORORIN exclude both rapid- and ultraslow-unloading regimes, constraining linker residence time to an intermediate range. WT and ΔSORORIN scsHi-C were reanalyzed from ref. (*21*). **(e)** SORORIN-dependent sequestration provides a similar degree of protection irrespective of unloading pathway details. Specifically we consider a model where SORORIN can bind linker cohesin irrespective of any other cofactor binding states prior to unloading, which specifies a three-state model. *Left:* Fraction of bound sister linkers over time in the full cofactor reaction network. *Right:* Fraction of bound sister linkers over time for the minimal three-state network. Transition rates were parameterized using the same FRAP data for SORORIN as the full model. At 16 h, SORORIN depletion increases the unloading rate by 2.5-fold in the full network and 2.1-fold in the three-state model. **(f)** Simulated linker loss following mitotic onset. Simulations were evolved under WT G_2_ maintenance kinetics until mitotic onset at 4 h (teal), when SORORIN rebinding was disabled to model its phosphorylation-dependent inhibition during mitosis (*42*, *43*). We then considered three scenarios. (i) If SORORIN provided protection through stable binding (10,000 s), inhibition of rebinding produced little increase in unloading rate. (ii) With the reference SORORIN residence time (100 s), bound SORORIN rapidly dissociates following inhibition of rebinding, resulting in a 2.5-fold increase in the unloading rate. (iii) An additional 30-fold stimulation of unloading allowed for removal of the majority of linkers within an amount of time compatible with prophase. Thus, while dynamic rather than stable SORORIN protection facilitates linker loss for mitosis, our analysis argues that additional pathways beyond simply blocking SORORIN rebinding operate *in vivo*.

**Figure S5.**
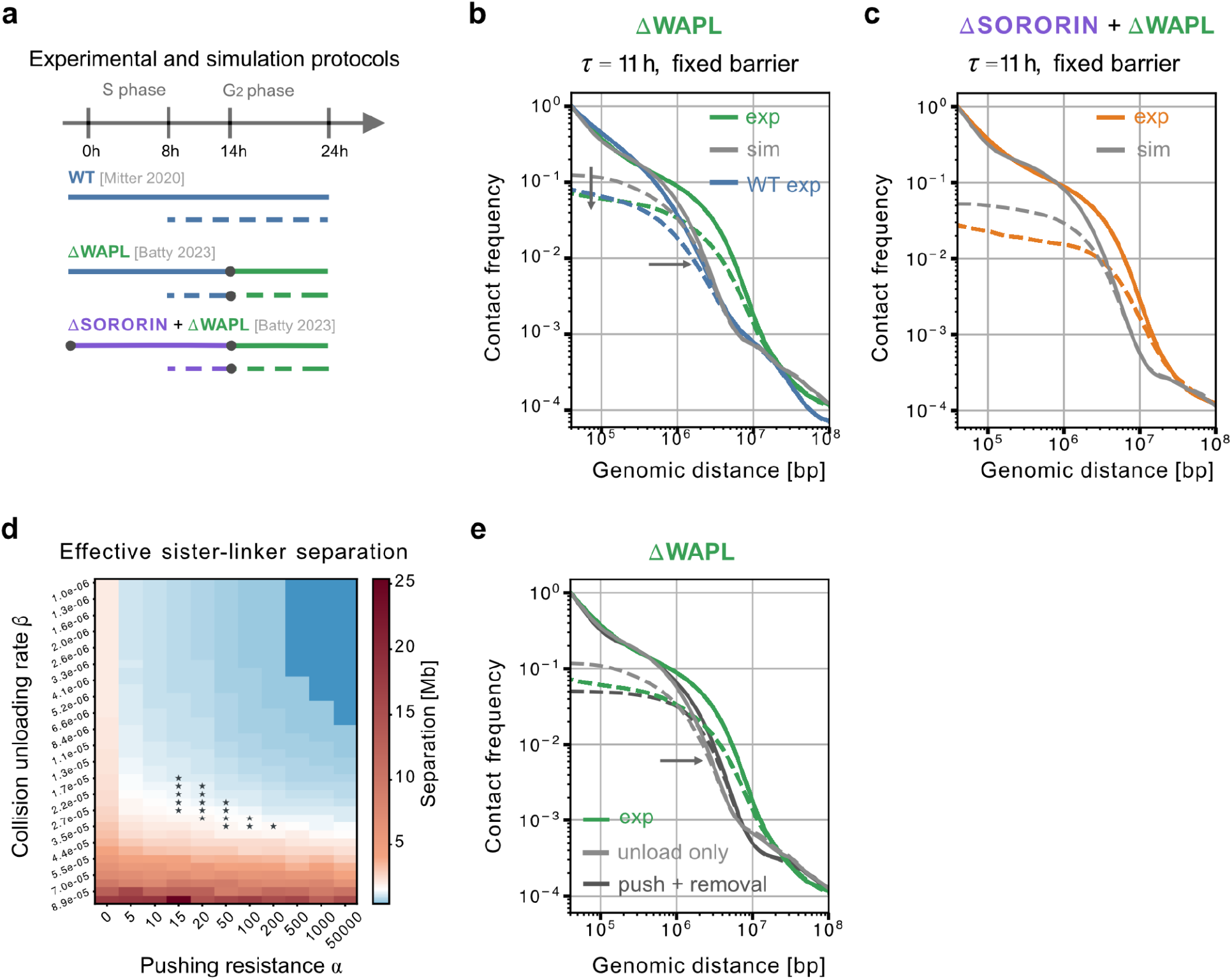
Extruder-linker encounter mechanisms are required to reproduce ΔWAPL P_trans_. **(a)** Experimental (solid) and simulation (dashed) protocols for time-resolved scsHi-C. WT cells were sampled at G_2_-16h (S24h). For ΔWAPL, WAPL was depleted at G_2_-6h; for ΔSORORIN+ΔWAPL, SORORIN was depleted before S phase entry and WAPL at G_2_-6h. Simulations were initialized from the inferred G_2_-0h configuration and propagated to G_2_-16h. WT scsHi-C data were reanalyzed from ref. (*21*); ΔWAPL and ΔSORORIN+ΔWAPL data were reanalyzed from ref. (*6*). **(b–c)** Experimental and simulated P_cis_ and P_trans_ with unloading alone under a fixed-barrier regime for ΔWAPL **(b)** and ΔSORORIN+ΔWAPL **(c)**. WT experimental curves are shown for reference. The fixed-barrier regime fails to reproduce both P_cis_ and P_trans_. **(d)** Effective sister-linker separation in ΔWAPL simulations across pushing resistance *α* and collision-mediated linker-removal *β*. Stars indicate parameter combinations consistent with constraints on linker separation, linker misalignment, and extruded loop size. **(e)** Experimental ΔWAPL scsHi-C compared with simulations incorporating unloading alone or both linker pushing and collision-mediated removal. Incorporating pushing and removal improves agreement with the experimental P_trans_ beyond that possible with unloading alone, but fails to reproduce P_cis_.

**Figure S6.**
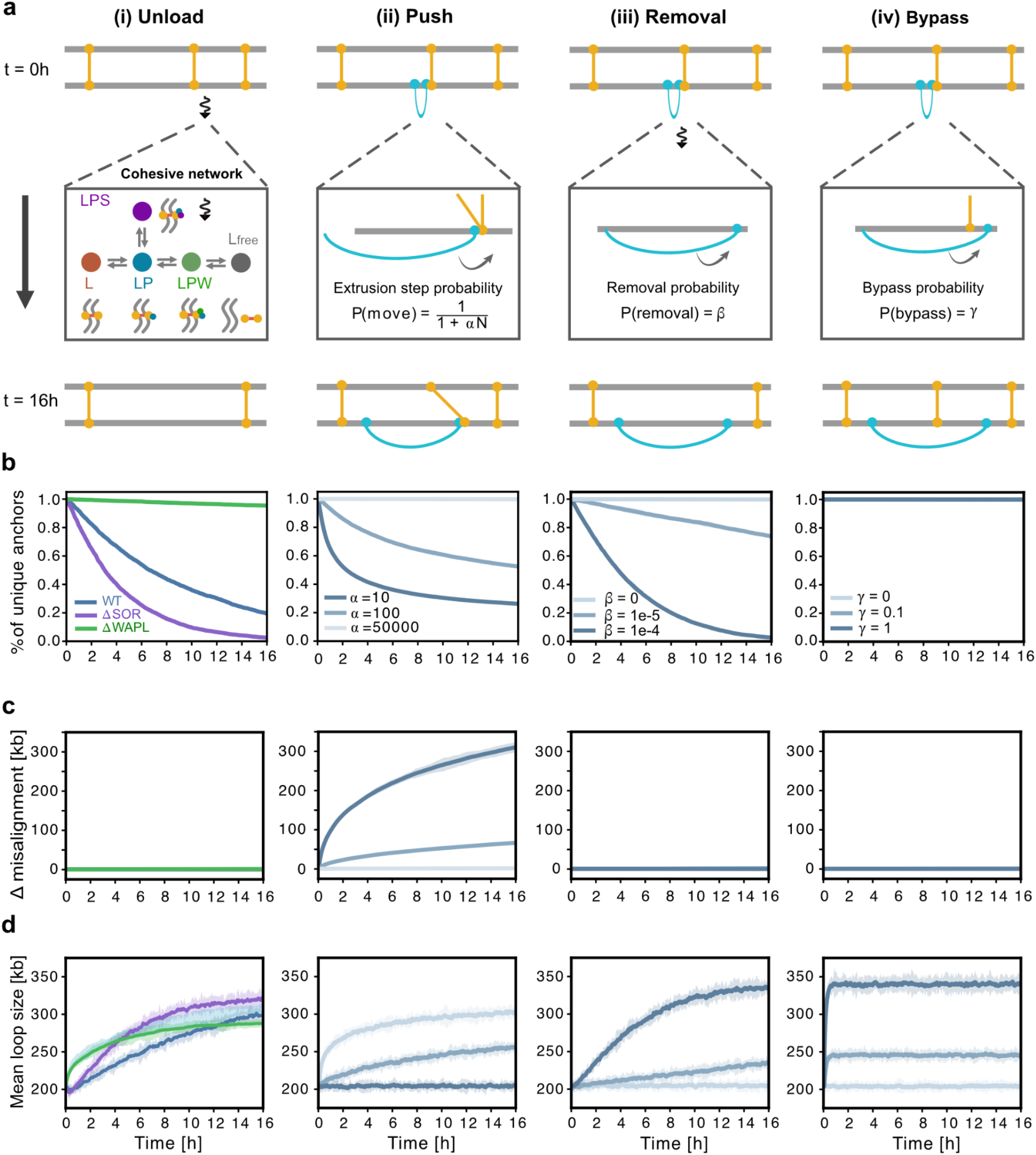
Multiple mechanisms can increase chromosome individualization during G_2_. **(a)** Lattice simulations of four mechanisms that alter the positioning of sister-linking and loop-extruding cohesins during G_2_: (i) linker unloading governed by the cofactor reaction network; (ii) linker pushing by extruders, with extrusion-step probability determined by pushing resistance *α* and the number of linkers pushed, N; (iii) collision-mediated linker removal with probability *β*; and (iv) linker bypass by extruders with probability *γ*. Simulations were propagated for 16 h from the indicated initial configurations. **(b)** Fraction of unique linker anchors remaining over time, normalized to 0 h. This metric captures loss of effective anchors through either linker removal or aggregation by pushing. **(c)** Mean linker misalignment over time, defined as the genomic displacement between paired anchor positions on the two sister chromatids. **(d)** Mean extruded loop size over time. Shaded regions indicate ±3 SD across 5 simulation replicates.

**Figure S7.**
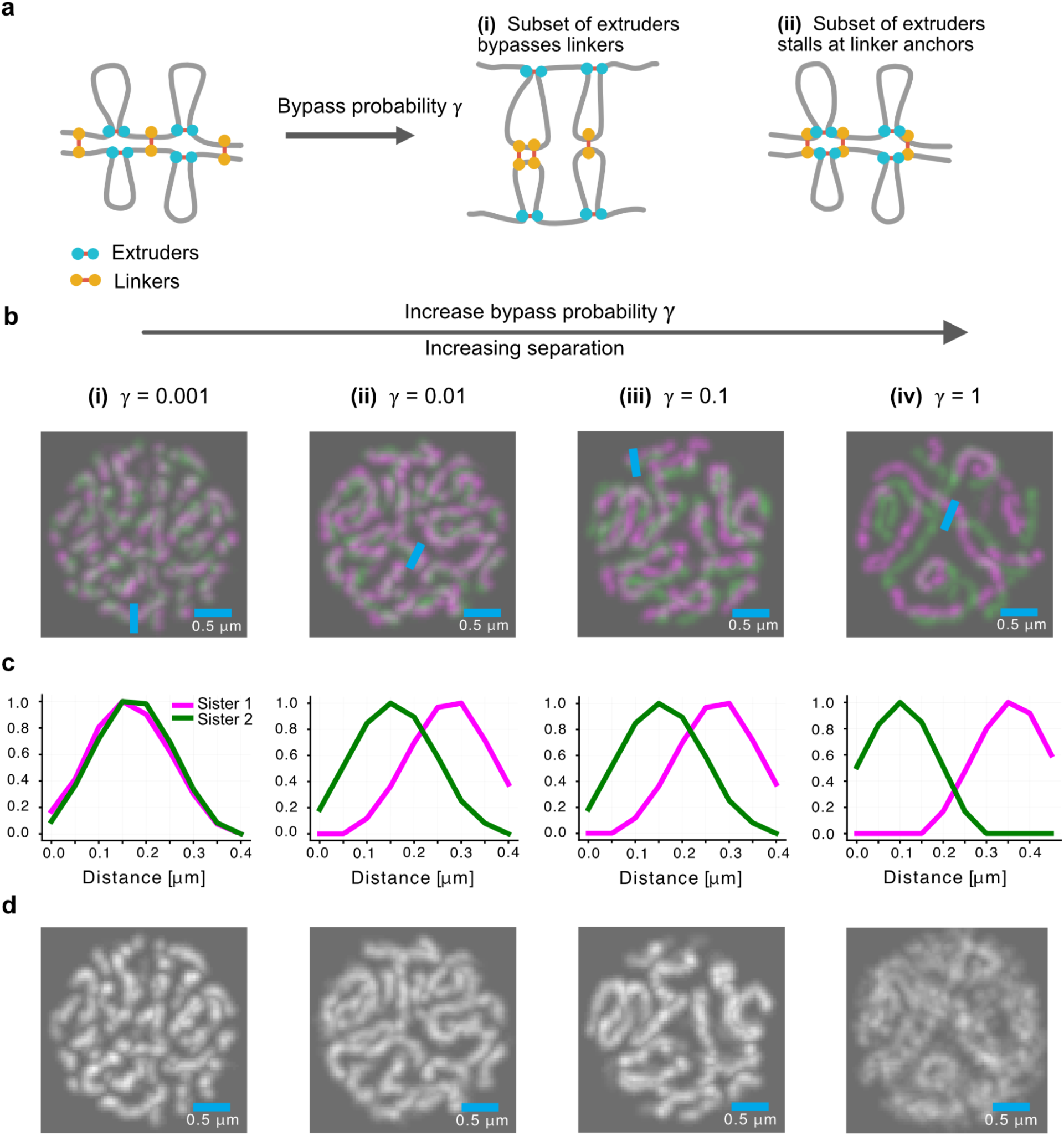
Chromosome morphology constrains extruder bypass to low probabilities. **(a)** Schematic of extruder-linker configurations generated by bypass. The rule of engagement defines how extruders interact with linkers. At G_2_-16h, a subset of linkers may be located within loops following bypass (i), whereas another subset may be stalled at loop anchors (ii). The relative abundance of these configurations is controlled by the bypass probability *γ* and determines chromosome morphology following ΔWAPL. **(b)** Representative simulated images of ΔWAPL chromosomes with increasing bypass probability (from *γ* = 0.001 to 1). Extruders on each sister chromatid are shown in magenta and green. Increasing bypass progressively separates the sister axes. **(c)** Normalized intensity profiles along the indicated line scans in **(b)**, showing increasing spatial separation of the two sister axes with bypass probability. **(d)** Corresponding grayscale simulated images showing all cohesins (linkers and extruders) combined.

**Figure S8.**
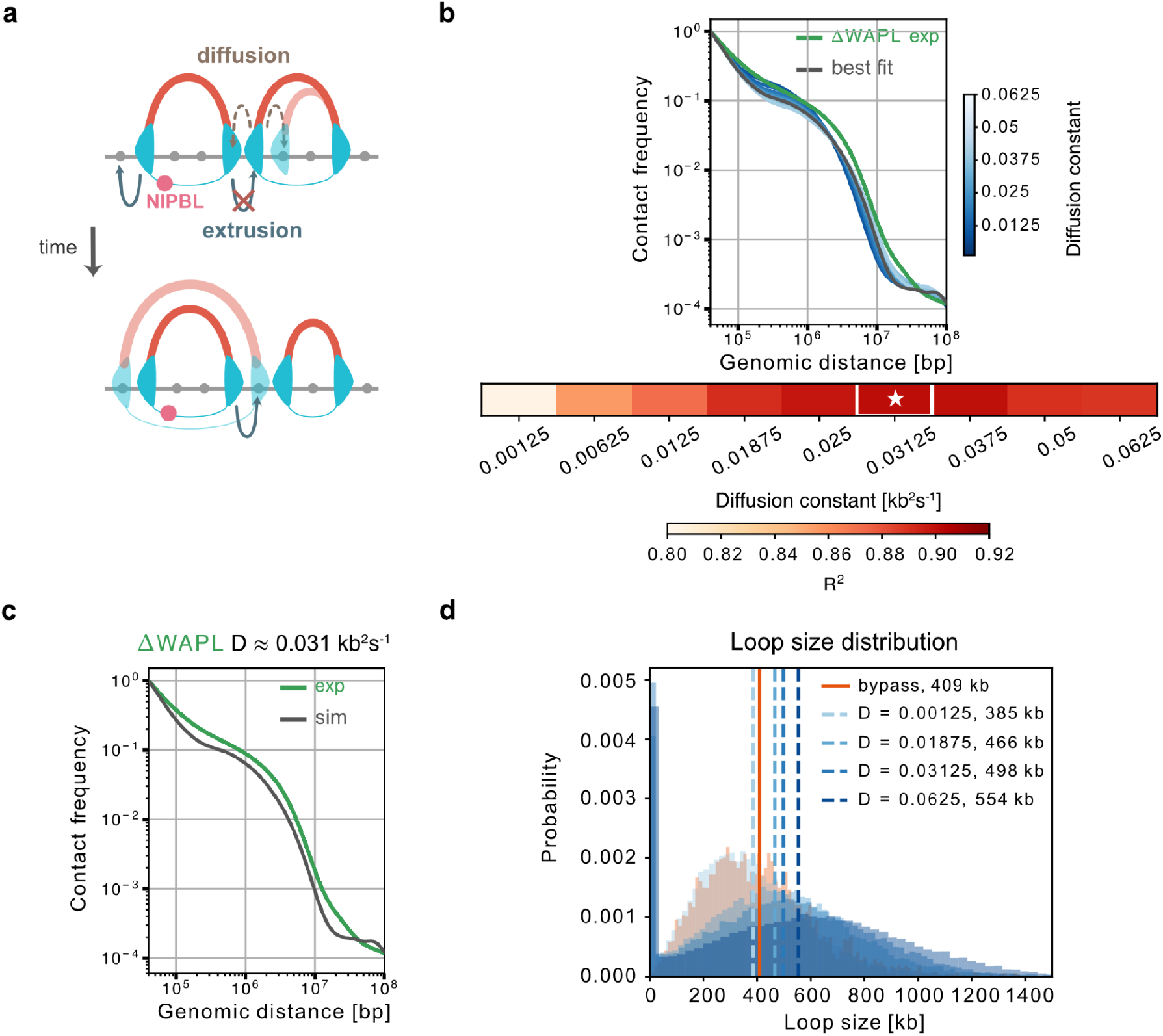
Extruder diffusion accounts for long-range cis contacts in ΔWAPL. **(a)** Schematic of how bi-directional extruder diffusion leads to a broader distribution of extruded loop sizes. Collisions between extruders block both active steps and diffusive steps. If an active extruder is collided with an inactive extruder, the inactive loop tends to shrink by diffusive steps, as the forward direction is blocked, and the active extruder moves into the vacated site. Over time this leads to additional growth of collided active loops. Thus diffusive motion in addition to loop extrusion provides a mechanism for additional loop expansion. **(b)** Dependence of simulated ΔWAPL P_cis_ on the extruder diffusion constant D. Top, experimental P_cis_ and simulations across different constants (blue), with the best-fitting simulation highlighted. Bottom, goodness-of-fit R^2^ to experimental ΔWAPL P_cis_ as function of D; the best-fitting value, D = 0.03125 kb^2^s^-1^, is indicated by the star. **(c)** Experimental and simulated ΔWAPL P_cis_ using the inferred diffusion constant. **(d)** Loop-size distributions generated by extruder diffusion across different D, with free bypass (*γ*= 1) shown for comparison. Increasing diffusion broadens the distribution and shifts its mean toward larger loop sizes. Dashed vertical lines indicate mean loop sizes. ΔWAPL scsHi-C data were reanalyzed from ref. (*6*).

**Figure S9.**
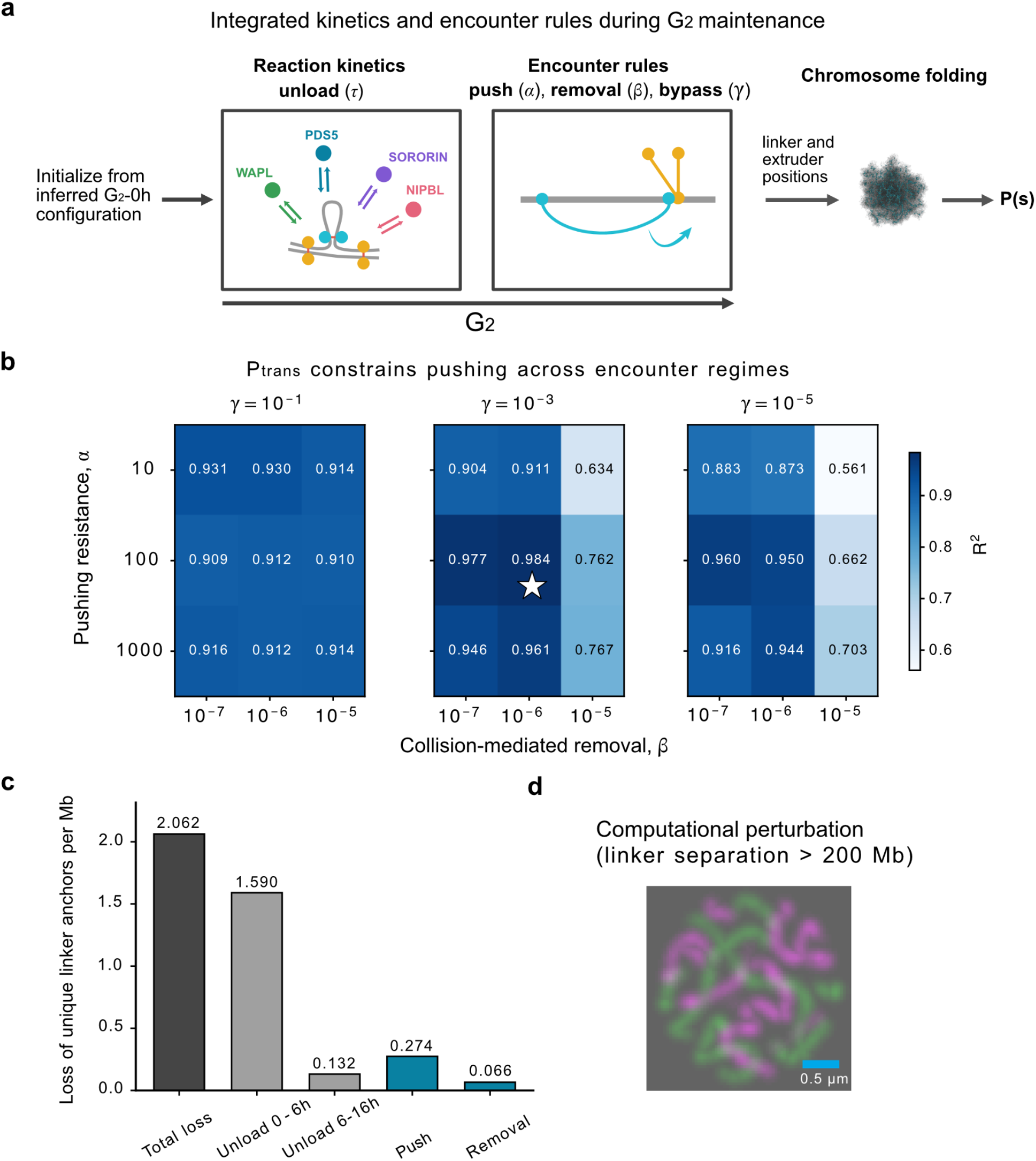
Integrated model identified by a joint sweep of pushing, removal and bypass. **(a)** Simulation workflow for the integrated model. Simulations were initialized from the inferred G_2_-0h (S8h) sister linker configuration and propagated through G_2_ with cofactor-dependent unloading and extruder-linker encounter rules acting jointly. Encounter rules were defined by pushing resistance *α*, collision-mediated removal probability *β* and bypass probability *γ*. The resulting linker and extruder positions were used as inputs for polymer simulations to calculate P(s). **(b)** Goodness of fit (R^2^) between experimental and simulated ΔWAPL P_trans_. Heatmaps show R^2^across pushing resistance *α*, collision-mediated removal *β* and bypass probability *γ*. Intermediate pushing provides better agreement with experimental P_trans_ than strong and weak pushing. The star indicates the inferred parameter combination. **(c)** Contribution of linker unloading, pushing, and collision-mediated removal to the loss of unique linker anchors over the 16 h G_2_ experimental window for the inferred integrated model. Pushing results in a 4.15-fold greater loss of linker anchors than collision-mediated removal, indicating that pushing is the dominant encounter mechanism reducing the effective number of linker anchors. Note that unloading mainly contributed to linker loss prior to 6 h and was greatly suppressed following WAPL depletion. Experimental ΔWAPL scsHi-C data used to calculate R^2^ were reanalyzed from ref. (*6*). **(d)** Computational perturbation of the inferred integrated ΔWAPL model testing an extreme regime with large linker separations. Extruders on the two sister chromatids are shown separately in magenta and green. Increasing linker separation beyond 200 Mb produces fully resolved sister chromatid axes, inconsistent with the experimentally observed ΔWAPL morphology.

**Figure S10.**
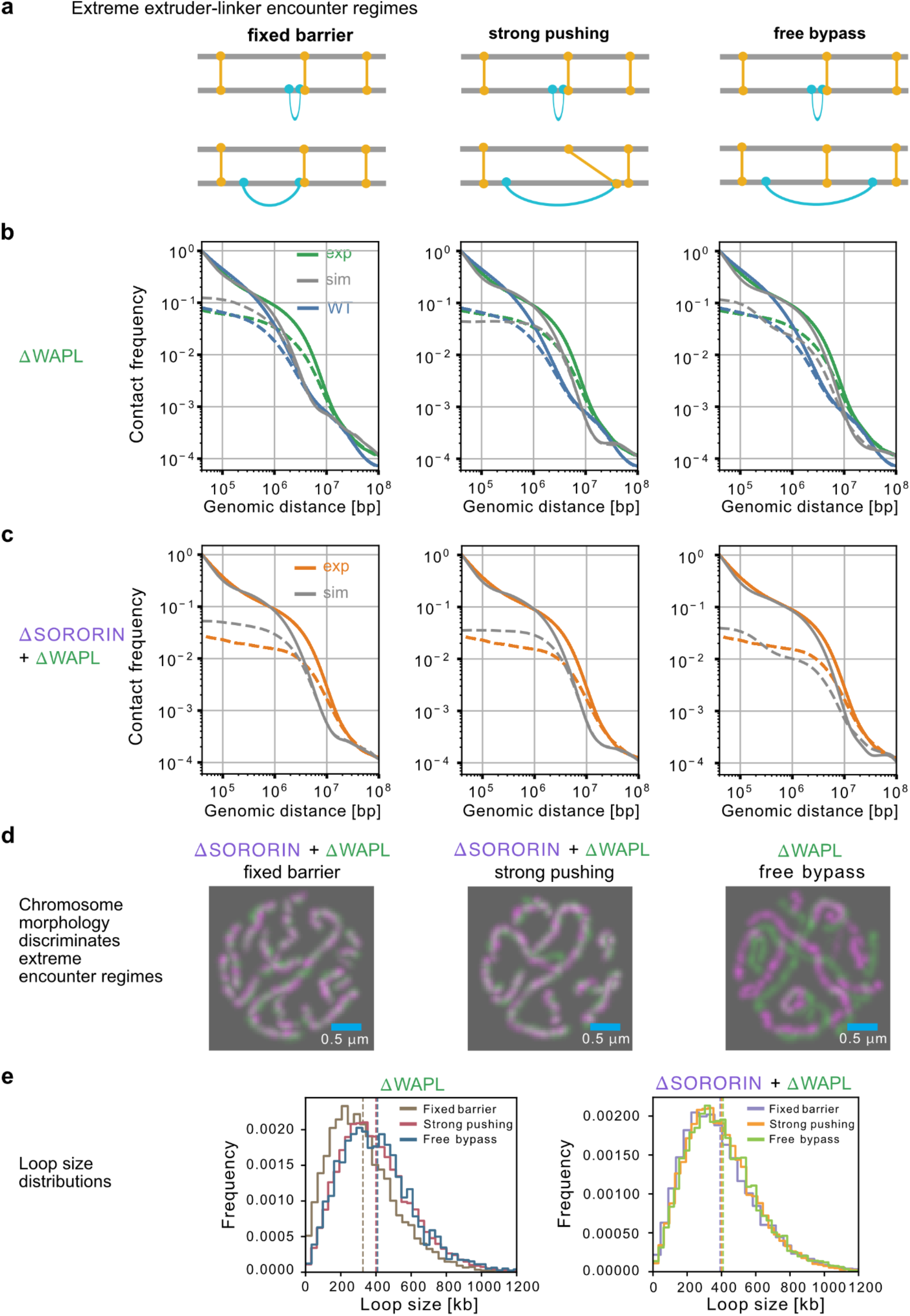
Extreme extruder-linker encounter regimes are inconsistent with WAPL-depleted chromosome organization. **(a)** Schematics of three extreme extruder-linker encounter regimes: fixed barriers, strong pushing, and free bypass. Fixed barriers (*α* = 50,000, *τ* = 12 h) prevent extruders from passing sister linkers; strong pushing (*α* = 5, *τ* = 12 h) allows extruders to displace linkers along the chromosome; and free bypass (*γ* = 1) allows extruders to pass linkers without displacing them. **(b–c)** Experimental and simulated contact-frequency curves under the three encounter regimes for ΔWAPL **(b)** and ΔSORORIN+ΔWAPL **(c)**. WT experimental curves are shown for reference. None of the extreme encounter regimes simultaneously reproduces P_cis_ and P_trans_ for both perturbations. **(d)** Simulated imaging provides an independent constraint on extruder-linker encounter rules. Fixed barriers and strong pushing fail to reproduce the separated sister axes observed in ΔSORORIN+ΔWAPL, whereas free bypass produces excessive sister-axis separation in ΔWAPL. **(e)** Loop size distributions for simulated ΔWAPL (left) and ΔSORORIN+ΔWAPL (right) under fixed barrier, strong pushing and free bypass regimes for extruder-linker encounters. Dashed vertical lines indicate the corresponding mean loop sizes. For ΔWAPL, mean loop sizes were 327 kb for fixed barriers, 400 kb for strong pushing, and 409 kb for free bypass. For ΔSORORIN+ΔWAPL, the corresponding mean loop sizes were 392, 400, and 410 kb, respectively. WT scsHi-C data were reanalyzed from ref. (*21*); ΔWAPL and ΔSORORIN+ΔWAPL data were reanalyzed from ref. (*6*).

**Figure S11.**
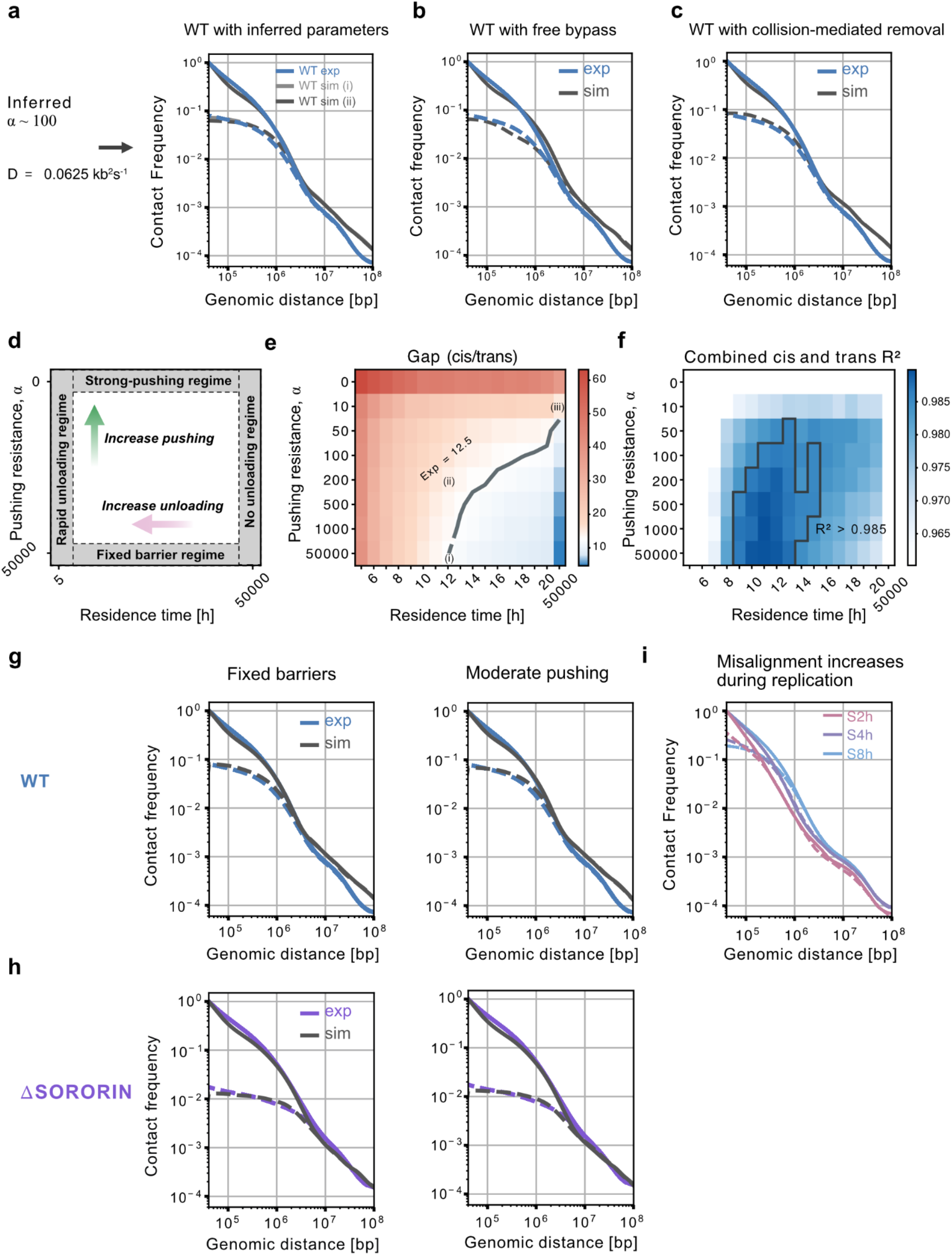
WT chromosome folding constrains but does not uniquely resolve cohesin encounter rules. **(a)** Experimental WT contact-frequency curves compared with simulations of (i) unloading alone (*α* = 50,000, *D* = 0) or (ii) unloading with pushing and diffusion using parameters inferred from ΔWAPL (*α* = 100, *D* = 0.03125 kb^2^s^-1^). Both simulations use a linker residence time of 11 h. The two parameter sets produce nearly indistinguishable P_cis_ and P_trans_ under WT conditions. **(b)** Allowing extruders to freely bypass linkers minimally alters simulated WT contact-frequency curves. **(c)** WT contact-frequency curves can also be reproduced by collision-mediated removal alone, illustrating the limited identifiability of encounter mechanisms under WT conditions. **(d)** Schematic showing how distinct pushing and unloading regimes emerge as a function of pushing resistance (*α*) and linker residence time (*τ*). **(e)** Simulated WT gap (P_cis_/P_trans_ at 40 kb) across pushing resistance *α* and linker residence time *τ*. The dashed line indicates the experimental WT gap (12.5). **(f)** Combined goodness of fit R² for P_cis_ and P_trans_ across pushing resistance *α* and linker residence time *τ*. The highlighted region indicates excellent fits ( R² > 0.985). Representative parameter combinations correspond to: (i) fixed barriers (*α* = 50,000, *τ* = 12 h); (ii) moderate pushing with unloading (*α* = 200, *τ* = 12 h); (iii) strong pushing without unloading (*α* = 10, *τ* = 50,000 h). **(g–h)** Experimental and simulated scsHi-C for WT **(g)** and ΔSORORIN **(h)** under fixed-barrier and moderate-pushing regimes. Both regimes closely reproduce the curves, indicating that WT and ΔSORORIN scsHi-C alone do not distinguish fixed barriers from moderate pushing. **(i)** Experimental scsHi-C during S-phase at 2, 4, and 8 h after release. The P_trans_ slope increases as replication progresses, consistent with increasing sister-linker misalignment. WT, ΔSORORIN and S-phase scsHi-C datasets were reanalyzed from ref. (*21*).

## Notes

### Competing Interest Statement

The authors have declared no competing interest.

