## Supplemental Information for "Chromosome folding encodes mechanisms of cohesion maintenance"

### Supplementary Information Content

Supplementary Table 1: Protein copy number, chromatin-bound fractions, and residence times for cohesin and its regulators 2

Supplementary Table 2: Literature estimates of stable chromatin-bound cohesin residence times 2

Supplementary Table 3: Summary of symbols used in the cohesin reaction networks 2

Supplementary Table 4: Reaction rates for the parameterized $G_2$ cohesin reaction network 2

Supplementary Table 5: Rules of engagement between loop-extruding and sister-linking cohesins 3

Supplementary Table 6: scsHi-C datasets and experimental conditions used in this study 3

Supplementary Table 7: Mechanistic inference from sister chromatid conformation 4

Supplementary Note 1: Kinetic formulation of the sister-linking cohesin reaction network 5

Supplementary Note 2: Approximate estimate of SORORIN-dependent protection 7

### Supplementary Information

### Supplementary Table 1

**Table S1.** Protein copy number, chromatin bound fractions, and residence times for cohesin and its regulators

| Protein | Copy number | Chromatin bound fraction | Chromatin residence time |
|---|---|---|---|
| **RAD21** | 284470 (average of refs. 33, 57) | Stable population: $1/3$ | 11 h (model inferred) |
| | | Dynamic population: $1/3$ | 822 s ($\mathrm{G_1}$; ref. 33)<br>600 s ($\mathrm{G_2}$; ref. 33) |
| **WAPL** | 69542 (average of refs. 33, 57) | 0.35 (ref. 10) | 45 s (ref. 10) |
| **PDS5** | 180615 (average of refs. 33, 57, 67) | 0.42 (ref. 10) | 72 s (ref. 10) |
| **SORORIN** | 79770 (average of refs. 33, 57) | 0.52 (ref. 10) | 100 s (ref. 10) |
| **NIPBL** | 119308 (average of refs. 33, 57) | 0.4 (ref. 59) | 72 s ($\mathrm{G_1}$; ref. 59)<br>80 s ($\mathrm{G_2}$; ref. 59) |

### Supplementary Table 2

**Table S2.** Literature estimates of stable chromatin-bound cohesin residence times

| Measurement | Residence time | Literature |
|---|---|---|
| FRAP with double-exponential fitting in HeLa cells in $\mathrm{G_2}$ | $6.3 \pm 3.7$ h<br>$8.6 \pm 4.1$ h | refs. 3, 33 |
| Immunoblotting of SMC3ac decay as an indirect proxy | 5 h | ref. 53, Supplementary Fig. S1a |
| iFRAP of STAG1 in $\mathrm{G_1}$ | 5 h | ref. 41, Fig. 2J |

### Supplementary Table 3

**Table S3.** Summary of symbols used in the cohesin reaction networks

| | |
|---|---|
| **Abbreviations** | |
| $P, W, N, S, R$ | PDS5, WAPL, NIPBL, SORORIN, and RAD21, respectively |
| $\mathrm{ext}, \mathrm{link}$ | Loop extruding and sister linking cohesin, respectively |
| **Reaction-network quantities** | |
| $\tau_X$ | Residence time of species $X$ |
| $N_X$ | Protein copy number of species $X$ |
| $F_X$ | Fraction of species $X$ bound to chromatin |
| $F_X^{\mathrm{ext}}$ | Fraction of all $X$ molecules associated with loop-extruding cohesin |
| $F_X^{\mathrm{link}}$ | Fraction of all $X$ molecules associated with sister-linking cohesin |

### Supplementary Table 4

**Table S4.** Reaction rates for the $\mathrm{G_2}$ cohesin network evaluated at the centroid of the feasible region, using a reference linker residence time of 10 h.

| Rate | Value | Unit |
|---|---|---|
| $k_{\mathrm{R,RP}}$ | $1.81 \times 10^{-7}$ | $\mathrm{molecule^{-1}s^{-1}}$ |
| $k_{\mathrm{RP,R}}$ | $1.57 \times 10^{-1}$ | $\mathrm{s^{-1}}$ |
| $k_{\mathrm{RP,RPW}}$ | $8.51 \times 10^{-7}$ | $\mathrm{molecule^{-1}s^{-1}}$ |
| $k_{\mathrm{RPW,RP}}$ | $2.19 \times 10^{-2}$ | $\mathrm{s^{-1}}$ |
| $k_{\mathrm{RP,RPS}}$ | $2.24 \times 10^{-6}$ | $\mathrm{molecule^{-1}s^{-1}}$ |
| $k_{\mathrm{RPS,RP}}$ | $1.00 \times 10^{-2}$ | $\mathrm{s^{-1}}$ |
| $k_{\mathrm{RPW,R_{free}}}$ | $3.15 \times 10^{-4}$ | $\mathrm{s^{-1}}$ |

### Supplementary Table 5

**Table S5.** Rules of engagement between loop-extruding and sister-linking cohesins

| Rule of engagement | Model definition |
|---|---|
| Pushing | Upon encounter, loop-extruding cohesins can push sister-linking cohesins along the chromosome. Pushing is controlled by the resistance parameter $\alpha$. |
| Fixed-barrier regime | Sister-linking cohesins act as fixed barriers to loop-extruding cohesins. In the pushing model, this corresponds to $\alpha \to \infty$. |
| Free-pushing regime | Loop-extruding cohesins push sister-linking cohesins without resistance. In the pushing model, this corresponds to $\alpha = 0$. |
| Bypass | Loop-extruding cohesins bypass sister-linking cohesins with probability $\gamma$ per extrusion step. |
| Free-bypass regime | Loop-extruding cohesins freely bypass sister-linking cohesins, corresponding to $\gamma = 1$. |
| Collision-mediated removal | Loop-extruding cohesins remove sister-linking cohesins upon encounter with probability $\beta$ per extrusion step. |

### Supplementary Table 6

**Table S6.** scsHi-C datasets and experimental conditions used in this study

| Dataset | Description | Literature | GEO |
|---|---|---|---|
| Unperturbed $G_2$-0h (S8h) | WT cells harvested 8 h after S phase release | ref. 21 | GSE152373 *Samples:* GSM4613673 |
| Unperturbed $G_2$-16h (WT) | WT cells harvested 24 h after S phase release | ref. 21 | GSE152373 *Samples:* GSM4613674 |
| $\Delta$SORORIN $G_2$-16h ($\Delta$SORORIN) | SORORIN depleted $\sim$1 h before S phase, achieving $\sim$97% depletion efficiency; cells harvested 24 h after S phase release | ref. 21 | GSE152373 *Samples:* GSM4613677 |
| $\Delta$WAPL $G_2$-16h ($\Delta$WAPL) | WAPL depleted 14 h after S phase release, achieving $\sim$99% depletion efficiency; cells harvested at 24 h | ref. 6 | GSE227694 *Samples:* GSM7105758–GSM7105763 |
| $\Delta$SORORIN+$\Delta$WAPL $G_2$-16h ($\Delta$SORORIN+$\Delta$WAPL) | SORORIN depleted before S phase and WAPL depleted 14 h after S phase release; cells harvested at 24 h | ref. 6 | GSE227694 *Samples:* GSM7105764–GSM7105767 |
| Unperturbed S2h | WT cells harvested 2 h after S phase release | ref. 21 | GSE152373 *Samples:* GSM4613671 |
| Unperturbed S4h | WT cells harvested 4 h after S phase release | ref. 21 | GSE152373 *Samples:* GSM4613672 |

### Supplementary Table 7: Mechanistic inference from sister chromatid conformation

**Table S7.** Experimental constraints on sister-chromatid maintenance mechanisms. Distinct chromosome-folding observables and perturbations constrain different components of the integrated maintenance model. Several mechanisms have little effect on WT chromosome folding and are therefore primarily resolved under perturbation (Fig. S11).

| Mechanism / model | Effect on cohesin organization | Key experimental constraint | Inference |
|---|---|---|---|
| WAPL-dependent unloading | Decreases the number of sister-linking cohesins during $G_2$ | WT and ΔSORORIN $P_{\mathrm{cis}}$ and $P_{\mathrm{trans}}$, particularly the gap (Fig. 3, Fig. S4) | Constrains linker residence time to an intermediate regime, excluding rapid and ultraslow unloading |
| Pushing | Displaces sister-linking cohesins, increasing effective linker separation and linker misalignment | ΔWAPL $P_{\mathrm{trans}}$ and the gap (Fig. 4, Figs. S5, S9, S10) | Moderate pushing is required under WAPL depletion but has little effect on $P(s)$ under WT conditions (Fig. S11) |
| Collision-mediated removal | Removes sister-linking cohesins upon encounters with extruders, increasing effective linker separation through loss of linker anchors | ΔWAPL $P_{\mathrm{trans}}$ and the gap (Figs. S5, S9) | Contributes to linker loss, but less strongly than pushing in the integrated model |
| Extruder bypass | Allows extruders to pass sister-linking cohesins, increasing loop size and promoting separation of sister-chromosome axes | ΔWAPL $P_{\mathrm{cis}}$ and simulated chromosome morphology (Figs. S7, S10) | High bypass probabilities are excluded by chromosome morphology, whereas bypass has little effect on WT $P(s)$ (Fig. S11) |
| Extruder diffusion | Redistributes extruders and broadens the loop-size distribution, increasing long-range cis contacts | Long-range ΔWAPL $P_{\mathrm{cis}}$ (Fig. S8) | Accounts for long-range cis contacts under WAPL depletion and contributes little under WT conditions (Fig. S11) |
| Integrated maintenance model | Combines the dominant contributions from cofactor-dependent unloading, pushing and extruder diffusion | ΔWAPL chromosome folding and morphology, with ΔSORORIN+ΔWAPL providing an additional perturbational test (Figs. S9, S10) | Reproduces multiple orthogonal features of WAPL-depleted chromosome organization |

#### Supplementary Note 1: Kinetic formulation of the sister-linking cohesin reaction network

The sister-linking cohesin reaction network comprises the following reactions:

$$L + P \underset{k_{\text{LP, L}}}{\overset{k_{\text{L,LP}}}{\rightleftharpoons}} LP \tag{S1}$$

$$LP + S \underset{k_{\text{LPS, LP}}}{\overset{k_{\text{LP,LPS}}}{\rightleftharpoons}} LPS, \tag{S2}$$

$$LP + W \underset{k_{\text{LPW, LP}}}{\overset{k_{\text{LP,LPW}}}{\rightleftharpoons}} LPW \tag{S3}$$

$$LPW \overset{k_{\text{LPW, free}}}{\longrightarrow} L_{\text{free}} + P + W \tag{S4}$$

Here, L denotes sister-linking cohesin, and the subscript "free" denotes unbound species. Mass-action kinetics give the following equations for the free species:

$$\frac{\partial [L]_{\text{free}}}{\partial t} = k_{\text{LPW, free}}[LPW] \tag{S5}$$

$$\frac{\partial [P]_{\text{free}}}{\partial t} = k_{\text{LPW, free}}[LPW] + k_{\text{LP, L}}[LP] - k_{\text{L, LP}}[L][P] \tag{S6}$$

$$\frac{\partial [W]_{\text{free}}}{\partial t} = k_{\text{LPW, free}}[LPW] + k_{\text{LPW, LP}}[LPW] - k_{\text{LP, LPW}}[LP][W] \tag{S7}$$

$$\frac{\partial [S]_{\text{free}}}{\partial t} = k_{\text{LPS, LP}}[LPS] - k_{\text{LP, LPS}}[LP][S] \tag{S8}$$

Consider a species $X$ that binds to chromatin with association rate $k_{\text{on}}$ and dissociation rate $k_{\text{off}}$:

$$[X]_{\text{free}} \underset{k_{\text{off}}}{\overset{k_{\text{on}}}{\rightleftharpoons}} [X]_{\text{bound}} \tag{S9}$$

The free population evolves as:

$$\frac{\partial [X]_{\text{free}}}{\partial t} = -k_{\text{on}}[X]_{\text{free}} + k_{\text{off}}[X]_{\text{bound}} \tag{S10}$$

At equilibrium, the bound fraction is:

$$f_X = \frac{[X]_{\text{bound}}}{[X]_{\text{bound}} + [X]_{\text{free}}} = \frac{k_{\text{on}}}{k_{\text{on}} + k_{\text{off}}} \tag{S11}$$

where $k_{\text{off}} = \frac{1}{\tau_X}$ and $\tau_X$ is the residence time of $X$.

We use the FRAP measurements of residence times and bound fractions to constrain the reaction rates. In the linker network derivation below, we omit the superscript “link”: $f_X$ and $N_X$ denote the bound fraction and copy number within the effective linker network pool, respectively. For each cofactor, this pool contains free molecules and molecules bound to sister-linking cohesin.

$$k_{\text{LP,LPW}}(f_P N_P - f_S N_S - f_W N_W) = \frac{f_W}{\tau_W(1 - f_W)} \tag{S12}$$

$$k_{\text{LPW,LP}} + k_{\text{LPW, free}} = \frac{1}{\tau_W} \tag{S13}$$

$$k_{\text{LP,LPS}}(f_P N_P - f_S N_S - f_W N_W) = \frac{f_S}{\tau_S(1-f_S)} \tag{S14}$$

$$k_{\text{LPS,LP}} = \frac{1}{\tau_S} \tag{S15}$$

$$k_{\text{L,LP}}(f_L N_L - f_P N_P) = \frac{f_P}{\tau_P(1-f_P)} \tag{S16}$$

$$k_{\text{LPW, free}} f_W N_W + k_{\text{LP,L}}(f_P N_P - f_S N_S - f_W N_W) = \frac{f_P N_P}{\tau_P} \tag{S17}$$

$$k_{\text{LPW, free}} N_W f_W = \frac{f_L N_L}{\tau_L} \tag{S18}$$

Solving these equations gives the rates:

$$k_{\text{L,LP}} = \frac{f_P}{\tau_P(1-f_P)(f_L N_L - f_P N_P)} \tag{S19}$$

$$k_{\text{LP,L}} = \frac{f_P N_P \tau_L - f_L N_L \tau_P}{\tau_P \tau_L (f_P N_P - f_S N_S - f_W N_W)} \tag{S20}$$

$$k_{\text{LP,LPS}} = \frac{f_S}{\tau_S(1-f_S)(f_P N_P - f_S N_S - f_W N_W)} \tag{S21}$$

$$k_{\text{LPS,LP}} = \frac{1}{\tau_S} \tag{S22}$$

$$k_{\text{LP,LPW}} = \frac{f_W}{\tau_W(1-f_W)(f_P N_P - f_S N_S - f_W N_W)} \tag{S23}$$

$$k_{\text{LPW,LP}} = \frac{-f_L N_L \tau_W + f_W N_W \tau_L}{f_W N_W \tau_L \tau_W} \tag{S24}$$

$$k_{\text{LPW, free}} = \frac{f_L N_L}{f_W N_W \tau_L} \tag{S25}$$

where $N_X$ is the copy number of species $X$, $\tau_X$ is the residence time of $X$.

### Supplementary Note 2: Approximate estimate of SORORIN-dependent protection

The linker unloading flux is:

$$\frac{dL_{\text{free}}}{dt} = k_{\text{off}}[L]_{\text{bound}} = \frac{1}{\tau_L}[L]_{\text{bound}} = k_{\text{off}}^{exit}[L]_{\text{exit}} \tag{S26}$$

Thus $\tau_L = \frac{[L]_{\text{bound}}}{[L]_{\text{exit}}} \frac{1}{k_{\text{off}}^{exit}}$.

Assuming that the intrinsic WAPL-dependent unloading rate is unchanged after SORORIN depletion, we obtain:

$$\tau_L^{\Delta SOR} \approx \frac{[L]_{\text{bound}} - [S]_{\text{bound}}}{[L]_{\text{exit}}} \frac{1}{k_{\text{off}}^{exit}} = \tau_L^{WT} - \tau_L^{WT} \frac{[S]_{\text{bound}}}{[L]_{\text{bound}}} \tag{S27}$$

thus,

$$\frac{\tau_L^{\Delta SOR}}{\tau_L^{WT}} \approx 1 - \frac{[S]_{\text{bound}}}{[L]_{\text{bound}}} \tag{S28}$$

Thus, the fraction reduction in linker residence time after SORORIN depletion is approximately equal to the fraction of linkers bound by SORORIN. Using the protein abundances and bound fractions in Table S1, we have $\frac{\tau_L^{\Delta SOR}}{\tau_L^{WT}} \approx 1 - \frac{f_S N_S}{f_L N_L} \approx 0.56$ (Table S1). Thus, under this rough estimate, linker residence time is reduced approximately twofold after SORORIN depletion. Using the scsHi-C S8h to estimate the bound RAD21, $\frac{\tau_L^{\Delta SOR}}{\tau_L^{WT}} \approx 1 - \frac{f_S N_S}{f_L N_L} \approx 1 - \frac{0.52 \times 79770}{19.539 * 1000/(800) * 2909} \approx 0.42$. The estimate likewise suggests approximately twofold protection by SORORIN.
